# The dissociative role of bursting and non-bursting neural activity in the oscillatory nature of functional brain networks

**DOI:** 10.1101/2022.11.02.514923

**Authors:** Alix Cordier, Alison Mary, Marc Vander Ghinst, Serge Goldman, Xavier De Tiège, Vincent Wens

## Abstract

The oscillatory nature of intrinsic brain networks is largely taken for granted in the systems neuroscience community. However, the hypothesis that brain rhythms—and by extension transient bursting oscillations—underlie functional networks has not been demonstrated *per se*. Electrophysiological measures of functional connectivity are indeed affected by the power bias, which may lead to artefactual observations of spectrally specific network couplings not genuinely driven by neural oscillations, bursting or not. We investigate this crucial question by introducing a unique combination of a rigorous mathematical analysis of the power bias in frequency-dependent amplitude connectivity with a neurobiologically informed model of cerebral background noise based on hidden Markov modeling of resting-state magnetoencephalography (MEG). We demonstrate that the power bias may be corrected by a suitable renormalization depending nonlinearly on the signal-to-noise ratio, with noise identified as non-bursting oscillations. Applying this correction preserves the spectral content of amplitude connectivity, definitely proving the importance of brain rhythms in intrinsic functional networks. Our demonstration highlights a dichotomy between spontaneous oscillatory bursts underlying network couplings and non-bursting oscillations acting as background noise but whose function remains unsettled.

**Significance statement:** Brain rhythms are paramount electrophysiological correlates of human cerebral activity as they coordinate neurons across distinct brain areas and establish neural synchronization. Spontaneous cortical oscillations, and particularly transient “bursts” of oscillations, also appear as the main electrophysiological subtrate of intrinsic functional brain networks. However, we argue that this oscillatory theory of brain networks, despite being widely accepted, should be requestioned due to a critical bias in electrophysiological measures of network connectivity. Here, we combined mathematical and neurobiological modeling techniques with magnetoencephalography recordings to set this theory on firm and rigorous grounds. Key to our analysis are scarcely studied non-bursting cortical oscillations. Although their precise function remains elusive, our results reveal that dissociating bursting and non-bursting oscillations is fundamental to the rigorous interpretation of electrophysiological network connectivity.

## 1. Introduction

A central notion in human neuroscience is that brain function is organized into intrinsic functional networks (Engel et al., 2013) sustained by spontaneously interacting neural assemblies (Deco and Corbetta, 2011; Fox and Raichle, 2007). Large-scale brain networks were initially mapped using resting-state functional magnetic resonance imaging (fMRI) (Biswal et al., 1995; Fox et al., 2005). Their electrophysiology and relation to brain rhythms were then investigated using resting-state magnetoencephalography (MEG) (Brookes et al., 2011; de Pasquale et al., 2010; Hipp et al., 2012). Electrophysiological networks turned out to be spectrally specific (Brookes et al., 2014, 2016; Tewarie et al., 2016; Wens et al., 2014a) and explainable in terms of sub-second transient “bursts” of brain rhythms (Baker et al., 2014; Coquelet et al., 2022; Hindriks and Tewarie, 2023; Seedat et al., 2020; Vidaurre et al., 2018b). Further studies also revealed their metastable, supra-second-scale dynamics underlying cross-network binding (Wens et al., 2019) and versatile network topology (de Pasquale et al., 2012, 2016; Della Penna et al., 2019) as well as their role in stimulus processing (Betti et al., 2018; Hawellek et al., 2013; Smith et al., 2009), task performance (O’Neill et al., 2015b; O’Neill et al., 2017; Quinn et al., 2018), learning and memory (Higgins et al., 2021; Mary et al., 2017; Roshchupkina et al., 2022; Van Dyck et al., 2021b), and several brain disorders (Brookes et al., 2018; Naeije et al., 2019; Puttaert et al., 2020; Sitnikova et al., 2018; Sjogard et al., 2020; Van Dyck et al., 2021a; Van Schependom et al., 2019).

The electrophysiology of intrinsic brain networks is strongly constrained by the amplitude envelope correlation (AEC) (O’Neill et al., 2015a; Sadaghiani et al., 2022; Siegel et al., 2012). The AEC is a functional connectivity measure that focuses on the oscillation amplitude of neural assemblies, tracks the spontaneous rise and fall of transient brain rhythms (Hari and Salmelin, 1997; van Ede et al., 2018), and quantifies to what extent these rhythms tend to burst simultaneously in distinct brain areas (Baker et al., 2014; Hindriks and Tewarie, 2023; Seedat et al., 2020). The co-occurrence of transient oscillatory bursts is actually thought to underlie electrophysiological network connectivity. This is different from the “communication-through-coherence” phenomenon (Fries, 2005), which instead depends on the precise time lag between these oscillations as measured by phase-based connectivity (Siegel et al., 2012). Crucially, these two theories of large-scale neural binding assume from the start that functional networks reflect interacting brain oscillations; yet this foundational assumption may be challenged. Conventional wisdom holds that signal amplitude or power artificially modulates connectivity estimation (Muthukumaraswamy and Singh, 2011; O’Neill et al., 2015b), an effect known as the power bias. The fact that electrophysiological functional networks are best delineated in the *α* (8–12 Hz) and the *β* (12–30 Hz) frequency bands (Brookes et al., 2011; Hipp et al., 2012; Siems et al., 2016; Wens et al., 2014a), precisely where resting-state signals exhibit their highest power (Hari and Salmelin, 1997), thus legitimately raises doubts on the oscillatory nature of functional connectivity. In a nutshell, connectivity could appear smaller in the lower (*δ, θ*) and higher (*γ*) frequency bands not because it is genuinely smaller but because electrophysiological recordings are noisier in these bands, leading to connectivity underestimation. So the possibility remains that intrinsic functional connectivity is broadband rather than carried by specific brain rhythms—a hypothesis first raised by Hipp and Siegel (2015). Experimental findings of broadband connectivity could lead to a paradigm shift in our conceptual understanding of the electrophysiology of functional networks (Cabral et al., 2014a; Deco et al., 2011).

Here, we sought to confirm or infirm the broadband connectivity hypothesis by directly disentangling the power bias from neurophysiological AEC in resting-state MEG recordings. We engineered a mathematically rigorous procedure that corrects for the power bias in spectrally resolved AEC. Critically, our procedure requires knowledge about the cerebral background processes that are not involved in amplitude coupling. This hinders *a priori* its applicability since un-mixing background and connectivity processes in resting-state data is a difficult, unsolved problem. We solved it here by incorporating a biologically informed model of amplitude coupling as oscillatory burst co-occurrence based on hidden Markov modeling (HMM) of MEG data. This unique combination of mathematical and neurobiological modeling allowed us to examine quantitatively how the power bias affects the connectivity spectrum of intrinsic functional networks and thereby demonstrate whether or not they reflect interacting neural oscillations.

## 2. Results

### The power bias reflects SNR-dependent connectivity under-estimation

We started our study with a theoretical analysis of the power bias in AEC. Figure 1(a–c) illustrates the most salient features of the resulting theory. (See Methods for a detailed formulation of the theory, and SI Theory for mathematical derivations.) We simulated pairs of synthetic electrophysiological signals mixing “connectivity processes” that generate a neural amplitude coupling and “cerebral background noise processes” that do not participate to this coupling. Simulations were performed at various levels of neural amplitude coupling and of signal-to-noise ratio (SNR), which turned out to be the main parameter controlling the power bias (see Methods). The relationship between AEC estimated from electrophysiological signals and the underlying neural amplitude coupling (AEC_0_) was linear with a slope that decreased when lowering the SNR (Fig. 1a). The slope was close to one at high SNR, indicating accurate amplitude coupling estimation when connectivity processes dominate over background noise (AEC ≈ AEC_0_); but then it decreased as the SNR got lower, revealing connectivity underestimation due to background noise. This SNR-dependent underestimation corresponds precisely to the power bias (Muthukumaraswamy and Singh, 2011; Hipp and Siegel, 2015).

**Figure 1.**
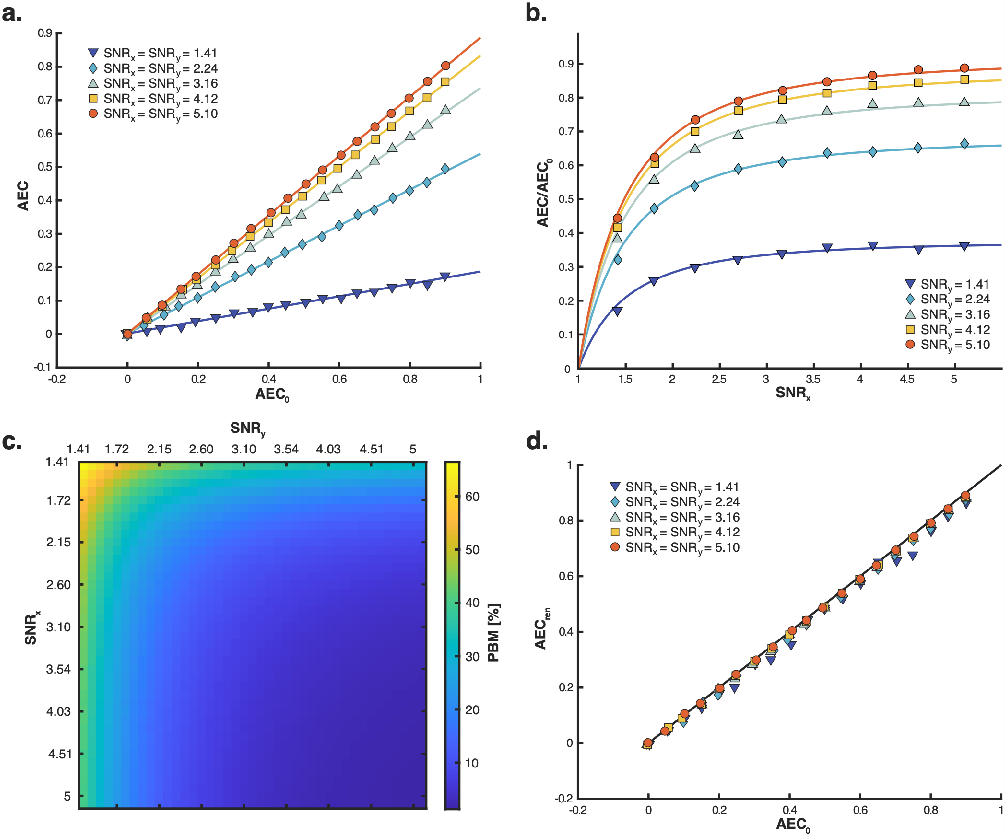
Power bias in amplitude correlation. Pairs of synthetic brain signals (*x, y*) were simulated at various SNR levels and neural amplitude couplings (AEC_0_). **(a)** Linear relationship between AEC estimation and AEC_0_. Linear regression curves are superimposed to data points. **(b)** Nonlinear SNR dependence of the power bias, illustrated by plotting the slope (AEC*/*AEC_0_ against the SNR. The superimposed model curves correspond to the nonlinearity (1 − 2 SNR^−2^)^1*/*2^ (see SI Theory). **(c)** Deviation of AEC estimates from the neural amplitude coupling measured as a percentage (power bias measure, PBM; see SI Methods) while systematically varying signal SNRs. **(d)** Connectivity corrected by renormalization (AEC_ren_) as a function of AEC_0_. The black diagonal line indicates perfect correction (AEC_ren_ = AEC_0_).

### A key characteristic of the power bias is its nonlinearity in the SNR

Figure 1b examines the slope mentioned above against the SNR of one signal. Connectivity raised sharply from zero at small SNR, where neural amplitude coupling is undetected because noise dominates over connectivity processes, and plateaued at large SNR where functional connectivity is detected—though still possibly underestimated due to noise in the other signal. The SNR of the second signal modulated this plateau in a similar nonlinear fashion. Unbiased estimation of neural connectivity was only reached when both SNRs were large enough. The combined effect of both SNR nonlinearities enables to assess the effect size of the AEC power bias quantitatively (Fig. 1c). Estimated AEC deviated from the simulated neural coupling AEC_*O*_ by 60% when the two signals exhibited a low SNR of 1.4, but this deviation decreased rapidly at higher SNRs to reach below 10% when both SNRs were above 3. We conclude that the AEC power bias becomes negligible once the SNR exceeds 3. (Of note, our general analysis of AEC disclosed two other sources of bias, i.e., background noise correlations and zero-lag synchronization processes; however, both of them were *a priori* expected to be subdominant compared to the power bias itself. See Methods and SI Theory for details, and SI Results for data-based evidence.)

### The power bias may be corrected by renormalizing AEC estimates

Figure 1(a) suggests that the underestimation effect of the power bias may be corrected by renormalizing AEC with the nonlinear slope factor illustrated in Fig. 1(b) (see Methods). Figure 1(d) provides proof-of-concept for this procedure using synthetic signals with time length and frequency content commensurate to the experimental MEG recordings analyzed below. The renormalization allowed to successfully recover the simulated neural amplitude coupling with high accuracy, even at the lowest SNRs (correction error = 0.4% at SNR = 1.4; *<* 0.1% for SNR *>* 3).

### Cerebral background noise may be identified as a non-bursting brain state

To apply our renormalization procedure to experimental MEG functional connectivity data, we first had to determine what the notion of “cerebral background noise” in our model represents in electrophysiological recordings. The problem was therefore to identify and isolate cerebral activity not involved in amplitude coupling. Given the neurobiological finding that AEC mostly reflects the coincident bursting of brain oscillations (Seedat et al., 2020), we tried and modeled cerebral background noise explicitly as non-bursting brain activity (Fig. 2). We identified non-bursting periods of MEG recordings as the low-amplitude state of a HMM applied locally to each brain node of the connectome (see Fig. 2a, for an illustration).

**Figure 2.**
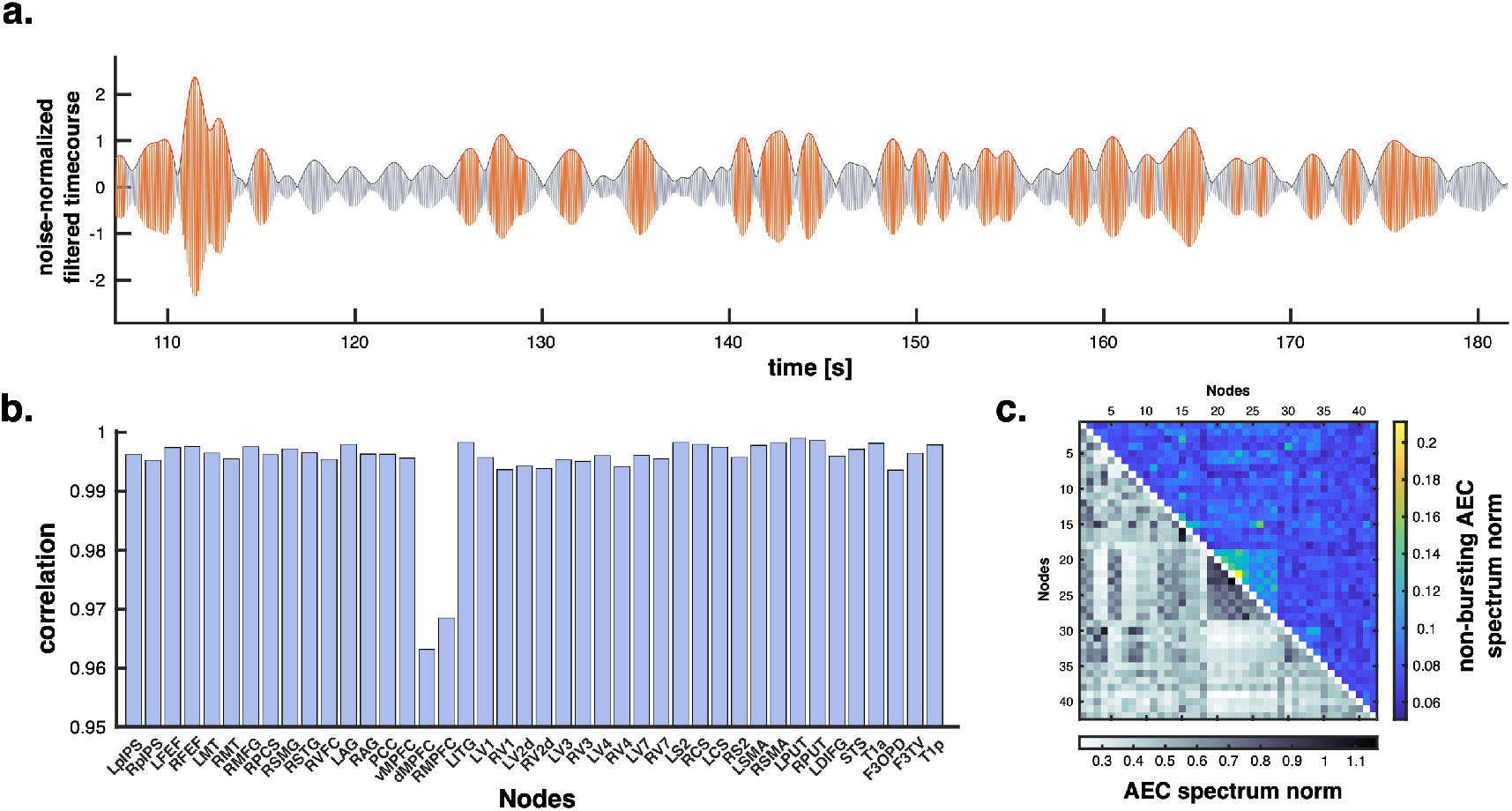
Cerebral background noise model based on non-bursting activity. **(a)** Illustration of the bursting (orange) and non-bursting (grey) states determined by the HMM applied to the MEG signal in the left sensorimotor network (node MNI coordinates, [− 42, − 26, 54] mm) filtered in the narrow [9.5, 10.5] Hz band. **(b)** Spectral similarity of non-bursting and bursting power spectra (see Fig. 3, bottom) using Pearson correlation at each of the 42 nodes of a whole-brain-covering connectome (de Pasquale et al., 2012). **(c)** Norm of non-bursting AEC spectra (upper right triangle) compared to bursting AEC spectra (lower left triangle). The smallness of non-bursting AEC norms indicates that non-bursting activity does not exhibit amplitude coupling (for explicit illustrations, see Fig. 3, top, black spectra).

To validate this approach, we started by analyzing interhemispheric AEC in three low-level brain networks (sensorimotor, SMN; auditory, AN; visual VN)—arguably the clearest hallmark of electrophysiological brain networks (Fig. 3, top). Their connectivity spectra exhibited typical peaks in the *α* and the *β* frequency bands (grey spectra), but they were flat for AEC corresponding to coincident non-bursting state activity in the two brain nodes (black). This was in stark contrast with the power spectra of non-bursting activity at the corresponding nodes (Fig. 3, bottom, green), which showed the same peaks than the oscillatory power spectra (i.e., SNR relative to measurement noise; blue spectra). The spectral similarity of non-bursting and bursting power spectra generalized across the whole brain (regularized Pearson correlation test; all *R*s *>* 0.96, *p*s *<* 10^−4^ controlling for the familywise error rate; Fig. 2b). Likewise, the spectral flatness of non-bursting AEC (Fig. 3, top black spectra) generalized across the whole connectome (Fig. 2c). We conclude that non-bursting activity is oscillatory but does not contribute to amplitude coupling. This means that non-bursting oscillations provide an adequate model of cerebral background noise to investigate the power bias in AEC. (See SI Theory for general conditions defining consistent noise models.)

**Figure 3.**
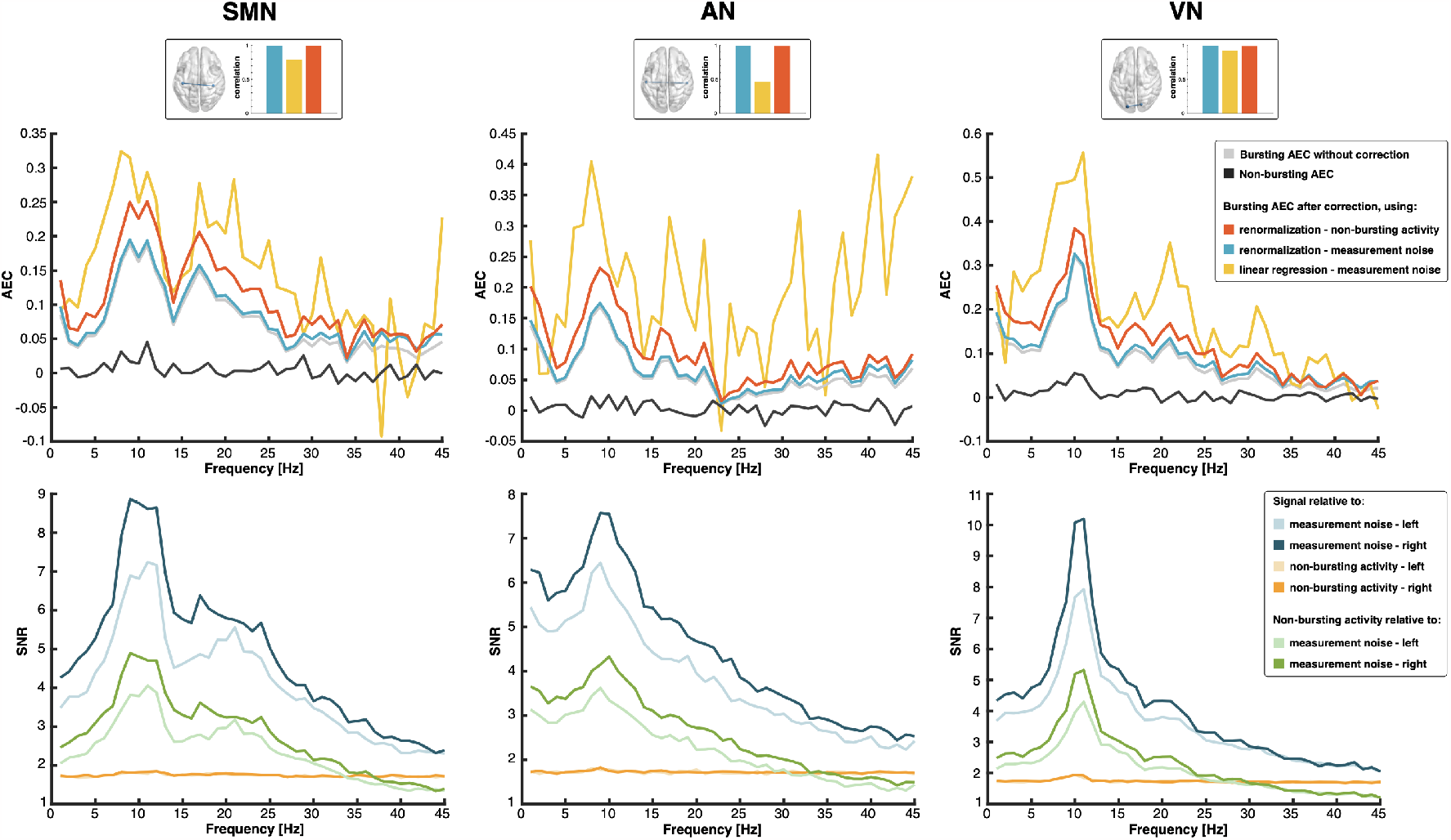
Power bias correction in interhemispheric amplitude correlation spectra. **Top**. Spectrally-resolved AEC is shown for three interhemispheric functional connections without correction (grey), after power bias correction based on AEC renormalization relative to cerebral background noise modeled as a non-bursting brain state (red) or to measurement noise (blue), and after linear regression (yellow). The AEC spectra of corresponding non-bursting activity are also shown (black; see also Fig. 2c for a summary measure of the spectral flatness of non-bursting AEC). Inserts show the connections on the MNI glass brain and compare the levels of spectral deformations (Pearson correlation between non-corrected and corrected spectra). **Bottom**. Spectrally-resolved SNR of MEG signals at the two nodes of each connection relative to cerebral background noise (orange) or to measurement noise (blue), and of non-bursting activity relative to measurement noise (green).

### The spectral content of interhemispheric connectivity networks is preserved by power bias correction

Figure 3 illustrates the fact that AEC spectral peaks (Fig. 3, top, grey) coincide with the presence of rhythmic brain activity at the corresponding nodes (Fig. 3, bottom, blue), which warrants a proper analysis of the power bias. The SNR relative to cerebral background noise (orange spectra) ranged from 1.7 to 2, for which simulations predicted AEC underestimation with effect size between PBM ≈ 5% and 30% (Fig. 1c at corresponding synthetic “gaussian” SNRs ranging from 2 to 3.5; see SI Theory for details on the link between experimental SNR and synthetic “gaussian” SNR). Accordingly, power bias correction levelled up AEC (Fig. 3, top, red spectra) with effect sizes in the expected range (PBM values: SMN, 18%; AN, 26%; VN, 17%). Crucially, the SNR relative to cerebral background noise still exhibited discernable *α*- and *β*-rhythm peaks reflecting oscillatory bursts, but the shape and peaks of AEC spectra were preserved after correction (regularized Pearson correlation test; SMN, *R* = 0.996; AN, *R* = 0.991; VN, *R* = 0.990). This confirms the physiological role of *α*- and *β*-bursts in primary functional networks.

We further assessed to what extent this conclusion depends on our model of cerebral background noise by reanalyzing power bias correction, only this time relative to measurement noise. This would amount to assume that the entirety of cerebral activity participates to amplitude coupling. Although detailed results differed, the end conclusion remained since spectral deformations were now minimal (Fig. 3, top, blue; PBM = 1% and *R*s *>* 0.995 for all three connections), consistently with the mixture of high (SNR = 2/”gaussian” SNR = 3, corresponding to predicted PBM below 10%) to very high SNR values relative to measurement noise (SNR = 10/”gaussian” SNR = 16 in the *α* band, corresponding to predicted PBM well below 1%; Fig. 3, bottom, blue spectra). On the other hand, the SNR nonlinearity of our model turned out to be key to draw our conclusion. Indeed, standard linear regression modeling proved inadequate for power bias correction (Fig. 3, top, yellow; see SI Results for more details).

### These conclusions generalize to the whole electrophysiological connectome

We then investigated AEC spectral deformations associated with the power bias systematically across the whole brain connectome (Fig. 4). Whatever the type of noise model considered in the correction, the shape of connectivity spectra was not deformed (regularized Pearson correlation, all *R*s *>* 0.898, *p*s *<* 1.3 10^−8^ after controlling for the connectome-level familywise error rate). The least similar spectra were located at a connection linking the visual (node MNI coordinates, [− 9, − 96, 13] mm) and the ventral-attentional networks ([41, 2, 50] mm) for power bias correction relative to cerebral background noise (Fig. 4, right; *R* = 0.977), and at a connection linking the auditory ([− 7, 9, 60] mm) and the control executive networks ([− 1, 10, 46] mm) for power bias correction relative to measurement noise (Fig. 4, left; *R* = 0.899). Inspection of these worst-case examples shows that spectral peaks in AEC connectivity are indeed all preserved after correction. We conclude that the power bias does not impact spectral features of intrinsic functional connectivity.

**Figure 4.**
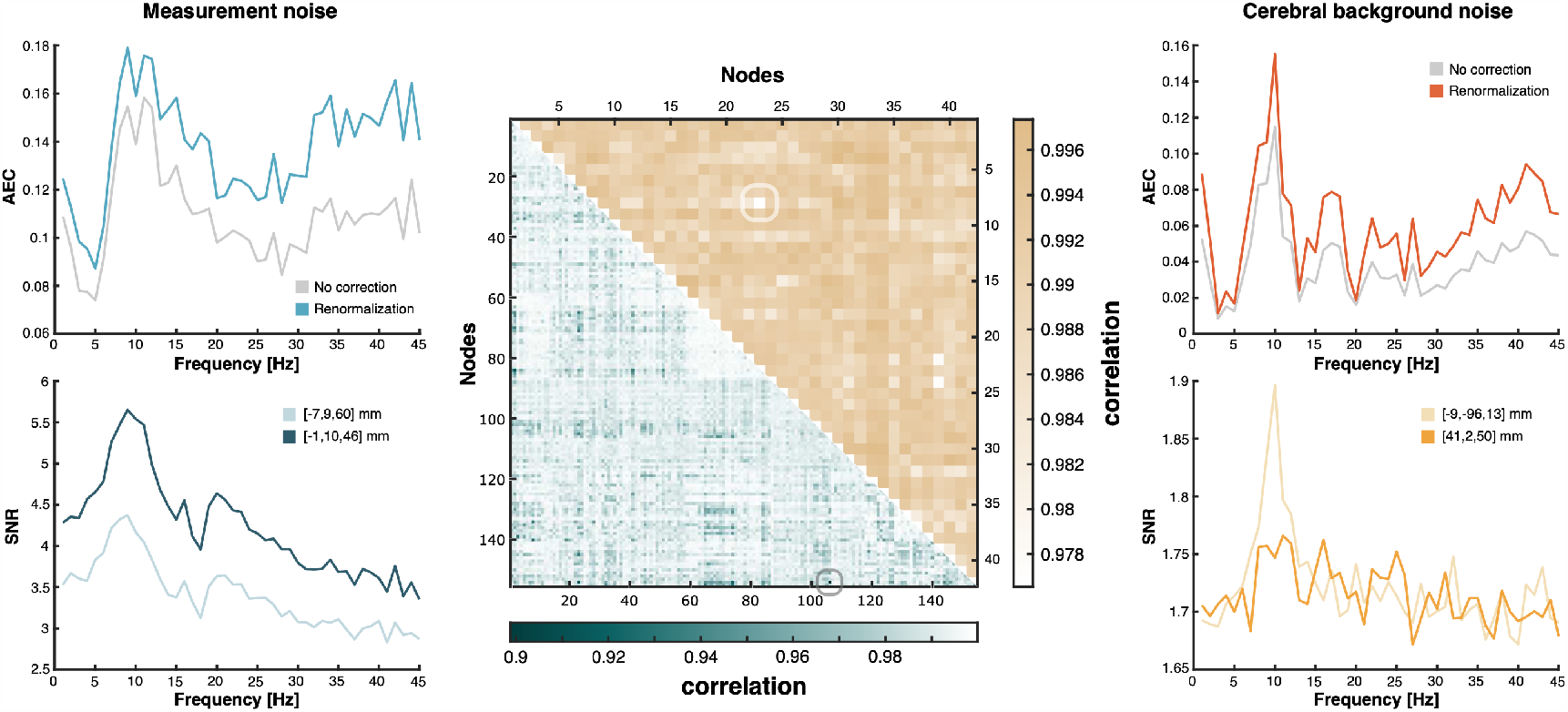
Power bias in amplitude correlation spectra of the connectome. **Middle**. The spectral deformation effect on AEC spectra due to power bias correction relative to cerebral background noise or to measurement noise was mapped in the whole-brain-covering connectome using a matrix of Pearson correlations between the non-corrected and corrected AEC spectra. A dense 155-node connectome (Della Penna et al., 2019) was used when using measurement noise (lower-left triangle), but for computational reasons a sparser, 42-node connectome (de Pasquale et al., 2012) was used with cerebral background noise (upper-right triangle). **Left**. Functional connectivity (top) and SNR (bottom) spectra of the connection with the worst spectral deformation when modeling noise as measurement noise. Its location in the connectome is highlighted in the corresponding correlation matrix. **Right**. Same as left, but modeling cerebral background noise as non-bursting brain activity.

## 3. Discussion

Combining a rigorous mathematical model of the power bias in frequency-dependent amplitude coupling with a biologically informed model of cerebral background noise as non-bursting brain oscillations, we demonstrated that the spectral content of resting-state MEG amplitude connectivity is preserved after power bias correction, despite fairly low SNRs relative to cerebral background noise. This provides strong empirical evidence that intrinsic functional network connectivity does reflect, from an electrophysiological standpoint, couplings among spontaneous brain rhythms.

### Intrinsic functional networks reflect interacting brain rhythms

The oscillatory nature of intrinsic functional connectivity at rest has been taken for granted since the first successful electrophysiological mappings of spectrally resolved brain networks with MEG (Brookes et al., 2011; Hipp et al., 2012), and was actually considered well before (Leopold et al., 2003). Still, the effect of the power bias (Muthukumaraswamy and Singh, 2011) was not controlled explicitly, leaving the door open that functional connectivity may not be specific to rhythmic activity (Hipp and Siegel, 2015). A direct consequence of our analysis is that intrinsic functional networks do correspond to interacting neural oscillations.

Our experimental evidence provides independent confirmation of the consensus on what are the neurophysiological mechanisms of intrinsic functional connectivity revealed by resting-state amplitude correlations (Cabral et al., 2014a; Deco et al., 2011). Large-scale neurocomputational models seeking to describe the underlying biophysics often share the same spirit (notwithstanding differences in their details). They start with local *γ*-band oscillations generated within each isolated cortical column, e.g., through feedback loops between excitatory and inhibitory neurons (Wilson and Cowan, 1972). These fast, local oscillations then interact via long-range excitatory synaptic couplings supported by the anatomical circuitry that connects remote populations. This results into a collective, brain-wide dynamics from which network-level brain rhythms unfold at lower frequencies, mostly in the slower *θ, α* and *β* bands due to conduction delays (Cabral et al., 2014b, 2017; Tewarie et al., 2019). In this framework, functional connectivity is thus an emergent dynamical phenomenon that can be described in terms of interacting neural oscillations. The significance of our data is to set this theoretical framework on firmer grounds. Large-scale neurocomputational models scarcely generate these local cortical oscillations from detailed neural membrane dynamics but rather use them as a starting point, either by modeling neural populations explicitly as oscillators with Kuramoto-like models (Breakspear et al., 2010; Cabral et al., 2011, 2014b), or as neural mass models with parameters chosen to exhibit a limit cycle or at least to stand at the brink of its bifurcation (Deco et al., 2009; Tewarie et al., 2019; Wilson and Cowan, 1972). Admittedly, population-level oscillations are generically observed in local, fully connected networks of spiking neurons (Gerstner et al., 1996); still other possibilities involving irregular, chaotic behavior have been described in local networks exhibiting excitation-inhibition balance (Brunel, 2000; Brunel and Wang, 2003; van Vreeswijk and Sompolinsky, 1996). Had power bias correction demonstrated a broad-band functional connectivity, the latter possibility could have been reconsidered. In fact, chaotic or stochastically driven models have been used in the context of fMRI connectivity (Deco et al., 2009; Honey et al., 2007), but to our knowledge their consequence at the level of MEG functional connectivity has not been investigated. Our result provides independent empirical data that such revision is, fortunately, not warranted. The conceptual framework based on the interplay between locally generated neural oscillations and large-scale delayed interactions, thus stands.

It is fair to mention at this point a methodological limitation of our MEG analysis framework, and of functional connectivity in general for that matter. Connectivity estimation provides only an indirect way to probe the neural interactions occurring in the brain and may be fraught with several interpretation issues, of which the power bias is but one example (Palva and Palva, 2012). In particular, functional connectivity measures may contain “ghost interactions” related to secondary spatial leakage effects and uncertainties in the exact location of nodes in the functional connectome (Colclough et al., 2015; Palva et al., 2018; Wens, 2015; Wens et al., 2015). This being said, most issues relate to spatial deformations rather than spectral deformations *per se*; e.g., they would lead to an artefactual spread of oscillatory connectivity across the connectome through ghost interactions. The persistence of the spectral content of amplitude connectivity after our power bias correction thus still provides evidence for the existence of a close relationship between interacting neural oscillations and intrinsic functional networks.

### Spontaneous brain activity contains non-bursting “background” oscillations unrelated to amplitude coupling

The next question is to know what kind of neural oscillation supports the functional binding of neural assemblies into intrinsic networks. Previous work established a key connection between amplitude connectivity at rest and transient oscillatory bursts (Seedat et al., 2020). These bursts emerge naturally in neuro-computational models near criticality, i.e., close to bifurcations of limit cycles, and indicate the presence of metastable brain oscillations (Freyer et al., 2011; Hindriks and Tewarie, 2023). On the other hand, the debate remains open on whether oscillatory bursts really differ from sustained brain rhythms, both from the electrophysiological and the functional perspectives (van Ede et al., 2018). Our biologically informed model of background noise in amplitude coupling brings further insight into this question.

While our study was not initially focused on the role of oscillatory bursts in functional brain networks, in the course of our analysis we used non-bursting activity to identify the part of neural signals disengaged from amplitude connectivity. We split our resting-state MEG data into a bursting phase characterized by transient increases of oscillatory amplitude, and a non-bursting phase characterized by lower levels of oscillatory amplitude. The non-bursting phase itself turned out to be oscillatory and to contain dominant, sustained rhythms in the *α* and the *β* frequency bands, but crucially it did not exhibit any amplitude correlation. This suggests that spontaneous brain activity contains two functionally distinct classes of spectrally similar neural oscillations: transient oscillatory bursts subtending amplitude connectivity (Seedat et al., 2020), and non-bursting background oscillations not involved in the process of amplitude coupling. The idea of considering brain rhythms as being composed of transient bursts has gained significant weight over the last years (Jones et al., 2016; van Ede et al., 2018) and its functional implications received a lot of attention, be it in relation to motor control (Bonaiuto et al., 2021; Feingold et al., 2015; Sherman et al., 2016), working memory (Higgins et al., 2021; Lundqvist et al., 2016), or resting-state functional connectivity (Baker et al., 2014; Coquelet et al., 2022; Hindriks and Tewarie, 2023; Seedat et al., 2020; Vidaurre et al., 2018b). On the other hand, the possibility and functional implications of non-bursting, possibly sustained, brain oscillations remain largely unexplored. Our usage of this concept was restricted here to the analysis of the power bias in amplitude connectivity, in which non-bursting oscillations are reduced to a mere “cerebral background noise”. Still, this terminology should not mislead us to think that they genuinely lack any functional relevance (quite analogously to how, in the past, spontaneous brain activity was discarded as unstructured brain noise in the preresting-state era; see, e.g., Deco and Corbetta, 2011). In fact, we envision that a larger field of research might develop around the study of non-bursting brain oscillations.

### Bursting oscillations provide a physiological link between oscillatory power and amplitude connectivity

Even though non-bursting background oscillations do not participate to amplitude coupling *per se*, they do affect amplitude connectivity estimation by playing the role of “cerebral background noise” in the power bias, i.e., noise with respect to the neural processes generating amplitude coupling. In fact, this allows to give a direct physiological interpretation to the power bias in amplitude correlation. It stands to reason that a brain activity exhibiting oscillatory bursts that are too rare or not ample enough (i.e., low SNR in our model) cannot be well discriminated from non-bursting oscillations and will thus show poor sensitivity to burst-dependent amplitude coupling (i.e., connectivity underestimation). Non-bursting oscillations could thus theoretically lead to connectivity spectral deformations, mostly outside the *α* and *β* frequency bands where oscillatory bursts are best identified (Seedat et al., 2020), and to a flattening of the connectivity spectrum after power bias correction. Our model attempted to properly disentangle the impact of non-bursting oscillations from the actual amplitude coupling generated by bursting oscillations. It did not reveal spectral deformations but rather demonstrated the presence of *α* and *β* spectral peaks in amplitude connectivity. This provides evidence that amplitude coupling is predominantly carried by *α* and *β* bursts rather than oscillatory bursts in other frequency bands (Seedat et al., 2020).

In this context, the spectral similarity between oscillatory power and functional connectivity appears to be physiological and not artefactual; both reflect the spectral content of their common denominator—the oscillatory bursts. That is why the broadband functional connectivity hypothesis of Hipp and Siegel (2015) may not be warranted after all. This hypothesis was based on the fairly reasonable assumption that spectral coincidences between measures of functional connectivity estimation and SNR must be the result of an artifact (i.e., the power bias). Our careful modeling of amplitude coupling suggests that they do reflect physiology, but it also highlighted how non-trivial it would be to reach this conclusion without the analysis tools developed here. Our model explains how spectral similarities between power (dominated by the contribution of non-bursting oscillations) and amplitude connectivity (driven by oscillatory bursts) largely reflect spectral similarities between non-bursting and bursting oscillations, but the latter coincidence remains unexplained at the moment. Following the abovementioned framework based on large-scale delayed interactions, a possible reason could be that non-bursting and bursting oscillations are generated by neural circuits with similar geometry and delays (leading to similar spectral content; Tewarie et al., 2019) but distinct stability properties (Freyer et al., 2011). However, pursuing this question goes beyond the reach of our analysis.

Another question that our analysis cannot elucidate is, what would be the possible function of this physiological relationship between oscillatory burst power and functional connectivity. Daffertshofer and van Wijk (2011) demonstrated explicitly how local oscillatory power may influence long-range synchronization through a purely neurodynamical mechanism. This was also illustrated by Tewarie et al. (2018) using simulated biophysical models. Interestingly, the latter work included discussion of the power bias in MEG connectivity; the authors argued that the relationship observed between power and phase couplings in MEG data is physiological because models do exhibit such relationship. Providing a direct proof of this claim would require extending our model to the context of phase connectivity, which represents a challenging avenue for future work (see SI Theory). This would be particularly valuable to try and dissociate the possibly distinct contributions of bursting and non-bursting oscillations to neural phase synchronization, and eventually to interpret rigorously findings of transient phase couplings based on time-embedded HMM of resting-state MEG recordings (which is partially confounded by power; Vidaurre et al., 2018b).

### Power bias correction by renormalization may become an important tool to interpret electrophysiological network connectivity at rest and in task

We investigated here specifically the oscillatory nature of intrinsic brain networks, but our power bias correction may also prove useful in cognitive neuroscience applications seeking to assess the impact of different tasks, behavioral conditions, or populations on functional connectivity (Muthukumaraswamy and Singh, 2011). Even though non-bursting oscillations do not affect the spectral features of amplitude connectivity, we observed underestimation effects that could reach substantial levels (up to PBM = 33% change after power bias correction; though this effect did not reach statistical significance, see SI Results). Our renormalization procedure allows precisely to infer what the burst-dependent amplitude correlation would be without the disturbances brought by non-bursting activity (which by the way makes our renormalized amplitude connectivity close in spirit to other burst-related connectivity measures such as burst coincidence, Seedat et al., 2020, or co-kurtosis, Hindriks and Tewarie, 2023). We conclude that power bias correction may be essential to properly interpret functional connectivity contrasts across behavioral or clinical conditions, especially in situations where brain rhythms are altered. Possibilities include, e.g., task-induced desynchronizations (Pfurtscheller and Lopes da Silva, 1999), pharmacological treatments (Leong et al., 2022), and brain pathologies (Babiloni et al., 2021). It is noteworthy that standard techniques based on within-condition power regression of functional connectivity data proved inadequate due to the inherent nonlinearity of the power bias in the SNR (see SI Results), which further emphasizes the importance of our model. This failure is the reason why in past publications, we could only rely on regression designs that suppress specifically the effect of between-condition power changes in corresponding connectivity differences without modeling its effect within each condition (Naeije et al., 2019; Sjogard et al., 2020; Van Dyck et al., 2021a).

In conclusion, our analysis of brain amplitude couplings allowed not only to prove the neural oscillation theory of intrinsic functional network connectivity, but also to disentangle the role of bursting and non-bursting oscillations. Further, it demonstrates the usefulness of moving away from generic linear modeling and towards theoretical developments that integrate rigorous mathematical analysis to neurobiologically informed models of brain dynamics.

## 4. Methods

### MEG data acquisition

We analyzed 31 healthy right-handed adult volunteers (16 females; mean age: 26.4 years, range: 19– 36 years; no history of neurologic or psychiatric disease) taken from a MEG resting-state dataset used in previous publications (Mary et al., 2015; Vander Ghinst et al., 2016), including functional network mapping (Wens et al., 2014b, 2019). Subjects signed a written informed consent and data usage was conform to the HUB–Hôpital Erasme Ethics Committee approval (References: P2011/054, P2012/049). Neuromagnetic activity was recorded (analog band-pass: 0.1–330 Hz, sampling frequency: 1 kHz) during 5 min at rest, while subjects gazed at a fixation point, using a 306-channel whole-scalp-covering MEG system (Vectorview Neuromag, MEGIN) inside a lightweight magnetically shielded room (Maxshield, MEGIN; see De Tiège et al., 2008 for details). Head movements were tracked with four head position indicator coils whose location relative to fiducials, was digitized beforehand along with the face and scalp (Fastrack Polhemus). A standard brain 3D T1-weighted magnetic resonance image (MRI) was also acquired using a 1.5T MRI scanner (Intera Philips) and co-registered manually with the head digitalization for individual head modeling and source reconstruction.

### Data processing

Environmental interferences and head movements were suppressed using the temporal extension of signal space separation (Maxfilter v2.2 with default parameters, MEGIN; Taulu et al., 2005) and physiological interferences (cardiac and ocular), with an independent component analysis (FastICA of MEG signals filtered between 0.5 and 45 Hz and projected on their 30 principal components; Vigario et al., 2000). These data were then decomposed spectrally into narrow frequency bands (band-pass filter centers: 1, 2, …, 45 Hz, band-width: 1 Hz) and source projected by minimum norm estimation on a 5-mm grid covering the MRI brain volume (see Wens et al., 2015 for implementational details).

Functional connectivity was estimated as the spectrally resolved AEC between pairs of source-projected MEG signals (Wens et al., 2014a; see also SI Theory), with geometric correction of spurious functional connectivity due to spatial leakage effects (Wens, 2015; Wens et al., 2015; Della Penna et al., 2019). Nodes of the interhemispheric connections between homologous cortices of primary networks (SMN, MNI coordinates: left node, [− 42, − 26, 54] mm, right node, [38, − 32, 48] mm; AN, left, [− 54, − 22, 10] mm, right, [52, − 24, 12] mm; VN, left, [− 20, − 86, 10] mm, right, [16, − 80, 26] mm) were obtained from previous references (de Pasquale et al., 2012, Hipp et al., 2012). The functional connectome was estimated as the AEC between the nodes of a dense, pointwise 155-node brain parcellation (Della Penna et al., 2019). Given the computational burden of the local HMM (see below), this connectome was subsampled to 42 nodes covering major resting-state networks (de Pasquale et al., 2012) whenever cerebral background noise was modeled as non-bursting activity. Since the geometric correction introduces small numerical asymmetries, any leakage-corrected AEC estimate between two source signals *x* and *y* was symmetrized by averaging it with the corresponding AEC between *y* and *x*.

### Power bias correction by renormalization

At the heart of our power bias correction is a mathematical model of amplitude coupling. We describe it fully in SI Theory. Here we summarize only the end result implemented in our MEG analysis pipeline. See SI Methods for an alternative (though ultimately inadequate) correction based on standard regression modeling.

Starting from an estimate of AEC between the narrow-band MEG signals *x* and leakage-corrected signal *y*, the neural part of amplitude correlation may be recovered as the renormalized estimate

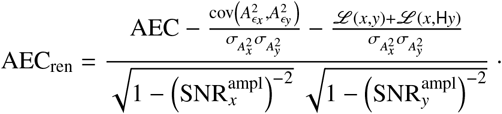

Like the original AEC, this quantity was eventually symmetrized over *x* and *y*. Here, *ϵ*_*x*_ and *ϵ*_*y*_ model the parts of signals *x* and *y* that do not contribute to amplitude coupling and thus play the role of “background noise” (see below), and H_·_ and *A* .denote the complex Hilbert transform and its amplitude (Hilbert envelope). The denominator corresponds to the renormalization correcting the power bias; it depends on an amplitude-specific version

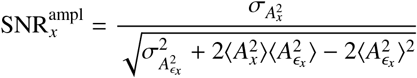

of the signal 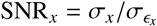 and similar expressions for signal *y* (see SI Theory for an explicit interpretation of 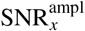). The second and third terms in the numerator further correct for possible noise correlations (although they were *a priori* expected to be subdominant by our definition of background noise, see SI Theory) and linear synchronization, the latter being controlled by the quantity

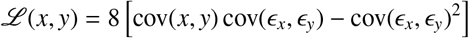

All temporal averages ⟨ · ⟩—including standard deviations *σ*_·_ and covariances cov (·, ·)—were estimated using all time points available in each individual recording. E.g., to model background noise as measurement noise, all statistics involving *ϵ*_*x*_ and*ϵ* _*y*_ were estimated from empty-room MEG signals at the brain locations corresponding to signals *x* and *y* (5 min; processed with same filters, source projection, and leakage correction than subjects’ recordings).

### Cerebral background noise model

To model background noise as non-bursting brain activity, we used a Gaussian two-state HMM (HMM-MAR toolbox; Vidaurre et al., 2016, 2018a) applied to the amplitude signal *A*_*x*_ (and leakage-corrected versions) at each brain node separately. Signal periods corresponding to non-bursting activity were inferred from the state showing the lowest mean amplitude with Viterbi decoding (Rabiner, 1989; Rezek and Roberts, 2005). Non-bursting power, nonbursting AEC, and all other statistics of *ϵ*_*x*_ and *ϵ*_*y*_ needed for power bias correction were computed by restricting time averages to coincident non-bursting periods both at signals *x* and leakage-corrected signals *y*. Non-bursting AEC was once again symmetrized over *x* and *y*. This enabled estimation of all the necessary statistical features of cerebral background noise, even though the HMM did not enable complete reconstruction of its time course simultaneously to the experimental recordings.

### Synthetic electrophysiological signals

We simulated neural connectivity processes as signal pairs *x*_0_, *y*_0_ initially generated as independent band-filtered gaussian white noises (8 − 12 Hz; 1 kHz sampling rate; 5 min) and then mixed nonlinearly together in order to introduce a “neural” amplitude correlation AEC_0_. Specifically, 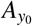 was replaced by 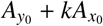 with

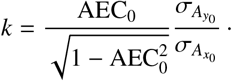

Background noise processes *ϵ*_*x*_, *ϵ*_*y*_ were simulated similarly but without amplitude correlation. Synthetic MEG signals were finally constructed by summing connectivity and noise processes, after independently rescaling them so as to fix SNR parameters. Simulations were run while controlling AEC_0_ and SNR parameters, using time length and number of repetitions matched to the size of our experimental resting-state dataset.

### Statistical procedures

The effect size of the power bias on amplitude correlation (power bias measure; PBM) was assessed as the relative difference between the estimated AEC and either the simulated coupling AEC_0_ or the corrected AEC_ren_ in experimental MEG data, measured globally over the whole frequency range (1–45 Hz). Spectral similarities in either oscillatory power or amplitude connectivity were assessed statistically using one-sided Pearson correlation tests. The null distribution of Fisher-transformed correlations *R* was Gaussian with zero mean and variance 1*/*(*n*− 3), the number *n* of spectral degrees of freedom being estimated from the cross-frequency covariance matrix of individual power or connectivity spectra (after averaging of the two spectra to be correlated). Significance was set to *p <* 0.05 with the familywise error rate controlled by Bonferroni correction for the number of independent nodes *ρ* estimated as the rank of the MEG forward model restricted to the nodes of the connectome (Wens et al., 2015) for power spectra, and for the number *ρ*(*ρ*− 1)*/*2 of independent connections in the connectome (Sjogard et al., 2019) for connectivity spectra. See SI Methods for full details.

## Author contributions

A.C. and V.W. designed study; A.M., M.V.G., and V.W. acquired data; A.C. and V.W. developed theory; A.C. contributed to analytical tools; A.C., X.D.T. and V.W. analysed data; A.C., A.M., M.V.G., S.G., X.D.T., and V.W. wrote and reviewed manuscript.

## Additional information

The authors declare that this research was conducted in the absence of any commercial or financial relationship that could be construed as potential conflict of interest.

## Data availability statement

Data will be made available upon request to the corresponding author and with the approval of institutional authorities (HUB–Hôpital Erasme and Université libre de Bruxelles).

## Code availability statement

Code will be made available upon request to the corresponding author.

## Acknowledgements

X.D.T. is Clinical Researcher at the Fonds de la Recherche Scientifique (F.R.S.FNRS, Brussels, Belgium). A.M. is Research Associate at the F.R.S.–FNRS. The authors acknowledge support from the F.R.S.–FNRS (research convention Excellence of Science EOS MEMODYN, 30446199). The MEG project at the HUB–Hôpital Erasme is financially supported by the Fonds Erasme (Research Convention “Les Voies du Savoir”, Brussels, Belgium).

## Supplemental Information

## Appendix A. SI Theory

We develop here in detail our mathematical theory of the power bias in spectrally resolved amplitude correlation. As a preliminary, we start with the simplest case of linear correlation, as it allows to illustrate clearly key theoretical features of the connectivity power bias, how a renormalization of functional connectivity enables to correct the bias, and what extra modeling steps are required for its implementation in practice.

### Appendix A.1. Warm-up: Linear correlation

#### Setup

The linear correlation *R* between two signals *x* = *x*(*t*) and *y* = *y*(*t*) is given by

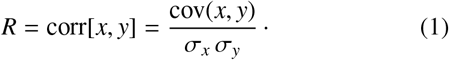

This corresponds to the covariance cov(*x, y*) = ⟨ *xy* ⟩ − ⟨ *x*⟩ ⟨ *y* ⟩of *x* and *y* suitably normalized by their standard deviation 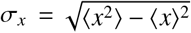 and 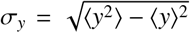. Brackets ⟨ ·⟩ denote time averaging. In our context, *x* and *y* represent experimental MEG signals reconstructed at two distinct brain locations. They can be decomposed as sums of the neural connectivity processes (respectively denoted hereafter as *x*_0_ and *y*_0_) that subtend the observed linear coupling, and of background noise (respectively *ϵ*_*x*_ and *ϵ*_*y*_) that may be of neural origin but do not participate to the generation of this coupling. Explicitly,

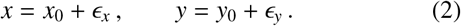

These simple-looking equations entail the hidden assumption that *x*_0_ and *y*_0_ are not mixed, as would be expected after source projection due to the spatial leakage effect (Wens, 2015). We shall keep this assumption since all our connectivity estimations included spatial leakage correction beforehand (Wens et al., 2015), notwithstanding possible remaining ghost interactions (Palva et al., 2018; see also main text for a discussion).

Our goal is to express the measured correlation (1) in terms of the neural coupling

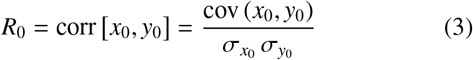

and the SNR estimates

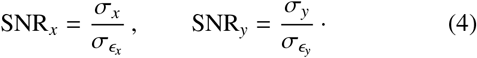

To keep the model as general as possible, we also leave the possibility of correlations *R*_noise_ = corr[*ϵ*_*x*_, *ϵ*_*y*_] among noises.

#### Power bias in linear connectivity

We demonstrate below that the above setup leads to the relationship

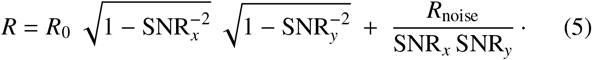

The correlation *R* can thus be decomposed into two distinct terms. The first encapsulates the power bias phenomenon and illustrates general features that we discuss now, momentarily neglecting noise correlations (i.e., we set *R*_noise_ = 0 in Eq. 5). First, the power bias leads to an underestimation of functional connectivity magnitude, |*R*| *<* |*R*_0_|. This effect is worst when noise dominates over connectivity processes (SNR_*x*_ »1 or SNR_*y*_ »1, so *R*≈ 0) but is negligible when connectivity processes dominate (SNR_*x*_ ≫ 1 and SNR_*y*_ ≈1, so *R* ≈*R*_0_). Second, this underestimation occurs through a SNR-dependent multiplicative factor. This is precisely what allows to disentangle the power bias from genuine functional connectivity, which can be recovered as a suitably renormalized version of the connectivity measure.

#### Power bias correction by renormalization

In the present case, and restoring the possibility of noise correlations, the *renormalized* correlation

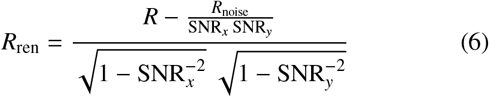

indeed allows to recover the neural correlation, *R*_ren_ = *R*_0_. Of notice, the renormalization factor (i.e., the denominator in Eq. 6) depends nonlinearly on the SNRs, indicating that conventional multiple regression modeling may not be able to efficiently correct the power bias. Numerically, the division entailed by the renormalization (6) must be well conditioned. This is not the case in the limit of low SNR (SNR_*x*_ ≈1 or SNR_*y*_≈ 1), for small estimation errors in the SNR estimates (4) then translate into large errors in the connectivity estimate (6). Therefore we predict that power bias correction will not be effective when noise dominates over the connectivity processes.

On top of the power bias itself, noise correlations *R*_noise_ contribute additively to functional connectivity estimates but their contribution is dampened at high SNR (second term in Eq. 5). Our strategy to model background noise in MEG data enforces that noise correlations vanish, so in theory *R*_noise_ = 0 (see defining property i below). In the context of MEG signals, persisting but spurious linear noise correlations are still bound to emerge in practice from linear mixing associated with source projection, although we can expect spatial leakage correction to dampen these correlations. So we can expect *a priori* that the power bias remains the dominating feature over noise correlations. Their effect can be eliminated by mere subtraction (see numerator in Eq. 6) alongside the power bias renormalization *per se*.

#### Modeling background noise

To make the correction procedure (6) applicable in practice, the SNR-dependent renormalization factor (as well as the noise correlation subtractive term) must be expressible in terms of parameters that are accessible to experimental data. We thus avoided formulating Eqs. (5) and in terms of, e.g., 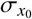 and 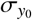 since the connectivity processes *x*_0_ and *y*_0_ are unknown. The noise processes *ϵ*_*x*_(*t*) and *ϵ*_*y*_(*t*) are also inaccessible, but second-order statistics (variances 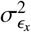 and 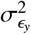 appearing through the SNR parameters, Eq. 4, and correlation *R*_noise_) are necessary. That is why our mathematical model must be supplemented with an explicit neurobiological model that isolates background noise processes and enables estimation of their statistics. Our broad strategy is as follows:

i. We define a model of *ϵ*_*x*_ and *ϵ*_*y*_ from signals devoid of the neural couplings of interest. In our analyses, we used either the “cerebral background noise” constructed from signal periods of coincident non-bursting activity in resting-state MEG recordings, or the “measurement noise” from empty-room MEG recordings (see main text). These model signals are not simultaneous with the experimental signals *x* and *y*, but they can be used nonetheless to estimate the required statistics. Note in particular that consistency of the model noises requires that their functional connectivity theoretically vanishes; e.g., we should expect *R*_noise_ ≈ 0 when investigating linear correlations (modulo spatial leakage effects).
ii. We further assume that the background noise processes *ϵ*_*x*_, *ϵ*_*y*_ contribute linearly to the experimental signals (2), and that they are independent of the neural connectivity processes *x*_0_, *y*_0_. This cannot be verified experimentally since by design the model signals are not simultaneous to the recordings *x, y*, although this is obviously true for measurement noise. For cerebral background noise, this amounts to genuinely assume that neural processes generating functional connectivity (e.g., oscillatory bursts; Seedat et al., 2020) and neural processes that do not participate to functional connectivity (e.g., non-bursting oscillations; see main text) are temporally independent.

#### Application to phase connectivity

It is noteworthy that the above considerations directly apply to the neuro-scientifically relevant phase connectivity measure known as coherence (Halliday et al., 1995), as it merely corresponds to a spectrally-resolved version of linear correlation. In this framework, results (5) and (6) hold with correlations (*R* and *R*_noise_) replaced by complex-valued coherency estimates and signal variances 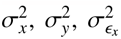, and 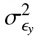 appearing in Eq. 4), by power spectral densities. The generalization to other phase connectivity measures such as the phase-locking value turns out to be much more challenging to handle. We do not pursue this here and consider in detail the case of amplitude connectivity relevant to the electrophysiological mapping of intrinsic brain networks.

#### Mathematical analysis: Derivation of Eq. (5)

We start by expanding the numerator in the right-hand side of Eq. (1),

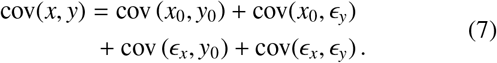

The two middle terms must vanish since connectivity processes *x*_0_, *y*_0_ and background noises *ϵ*_*x*_, *ϵ*_*y*_ are assumed to be independent, leaving cov(*x, y*) = cov (*x*_0_, *y*_0_) + cov(*E*_*x*_, *E*_*y*_). Similarly, in the denominator

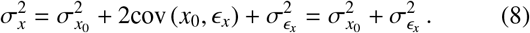

Combined with our definition (4) of the SNR, this equation can be recast as

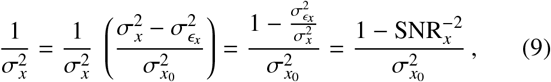

the factor between parentheses being equal to one. A similar equation holds for the signal *y*. Plugging these results back into Eq. (1) and using Eq. (4) once more leads to the sought Eq. (5).

### Appendix A.2. Amplitude correlation

#### Setup

Assessing the simultaneity of the rise and fall of transient neural rhythms in two brain areas requires first to isolate the amplitude time course of these rhythms and then to assess their temporal covariation. Isolation of rhythmic activity itself in MEG signals can be performed by band-pass filtering, so we will assume hereafter that *x* and *y* correspond to sufficiently narrow-band signals. In this context, we may interpret the signals *x*_0_ and *y*_0_ in Eq. (2) as local neural rhythmic activity that generates their functional coupling, and *ϵ*_*x*_ and *ϵ*_*y*_ as the band-filtered part of the rest of local neural activity not involved in this functional coupling (possibly wideband, though our results suggest that they contain sustained non-bursting oscillations; see main text) along with measurement noise.

Time-varying oscillatory amplitudes can then be conveniently extracted using the Hilbert transform, which factorizes any band-limited signal, say *x*(*t*), as 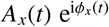, where *A*_*x*_ denotes its amplitude time course (a.k.a. Hilbert envelope) and *ϕ*_*x*_ its instantaneous phase time course. This factorization applies to both connectivity and noise processes, so the relation *x* = *x*_0_ + *ϵ*_*x*_ becomes

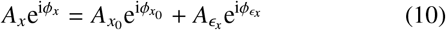

and similarly for the signal *y*. The mathematics of Hilbert transformation needed to develop our theory is briefly reviewed below.

In this framework, the co-occurrence of transient brain rhythms in two brain signals *x* and *y* translates into the temporal dependence of their two oscillatory amplitudes *A*_*x*_ and *A*_*y*_, which is conventionally measured as their correlation, i.e., AEC = corr[*A*_*x*_, *A*_*y*_]. Our goal is to express this functional connectivity measure in terms of the neural 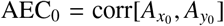 involving only oscillatory amplitudes 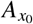 and 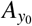 generating the coupling, and of a version of the SNR pertaining to oscillatory amplitudes that will be defined below.

#### Amplitude vs. power correlation

The major technical difficulty in this endeavor is that *A*_*x*_ is a nonlinear function of 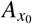 (see Eq. 30 below), preventing us to follow the strategy used in Appendix A.1 for linear correlation. Fortunately, it turns out that the correlation of amplitudes in the AEC can be replaced, to an extremely good approximation, by the correlation of amplitudes squared (Wens, 2015), so

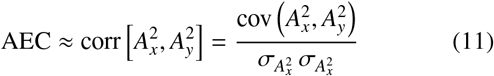

and similarly for AEC_0_. The correlation of amplitudes squared is actually closely related to the band-limited power correlation used by de Pasquale and colleagues (de Pasquale et al., 2010; Della Penna et al., 2019), which has been observed not only to yield the same functional brain networks than AEC but also quantitatively very similar functional connectivity estimates (Sjogard et al., 2019). The mathematical derivation of the approximation (11) is derived in Appendix E of Wens (2015). It is noteworthy that this approximation could have been avoided altogether, had we measured intrinsic functional connectivity with power correlation from the start; we did not do so here to keep in line with a large portion of the literature on MEG amplitude connectivity.

#### Power bias in amplitude correlation

Using Eq. (11) as starting point, we show below how to generalize Eq. (5) for linear correlations to the case of narrow-band AEC. The main result of this analysis reads

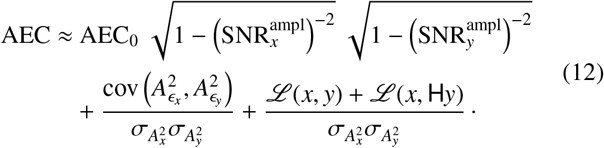

The first term embodies the power bias in AEC. It is formally similar to that in linear correlation (Eq. 5) except that it involves an amplitude-specific version of the SNR defined as

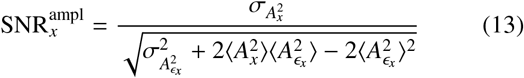

and similarly for *A*_*y*_. To get some intuition on this unusual SNR measure, we plot in Fig. S1 this quantity 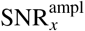 as a function of the standard SNR_*x*_ (Eq. 4) for the synthetic electrophysiological signals used to explore our theory numerically (see main text). In this case, the rel<u>a</u>tion appeared linear, with a slope estimated as 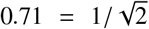 by linear regression, so the nonlinearity factor 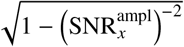 may be simplified to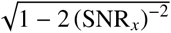 (see caption of Fig. 1). It turns out that this simplification emerged in this plot only because our synthetic signals were close to being gaussian (see the last paragraph of this Appendix) but it does not hold in general for non-gaussian signals such as experimental MEG recordings. For that reason, the predicted power bias effect sizes reported in Fig. 1(c) expressed in terms of the “gaussian” SNR may not be directly interpreted in terms of the SNR of MEG signals. Since Eq. (12) demonstrates that the power bias in AEC only depends on the SNR through the amplitude-specific version (13), the correct way to read off predictions from Fig. 1(c) is to estimate 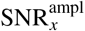 from experimental MEG recordings, artificially convert them into “gaussian” SNR according to 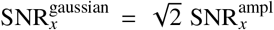 (Fig. S1), and use these values in Fig. 1(c).

**Figure S1:**
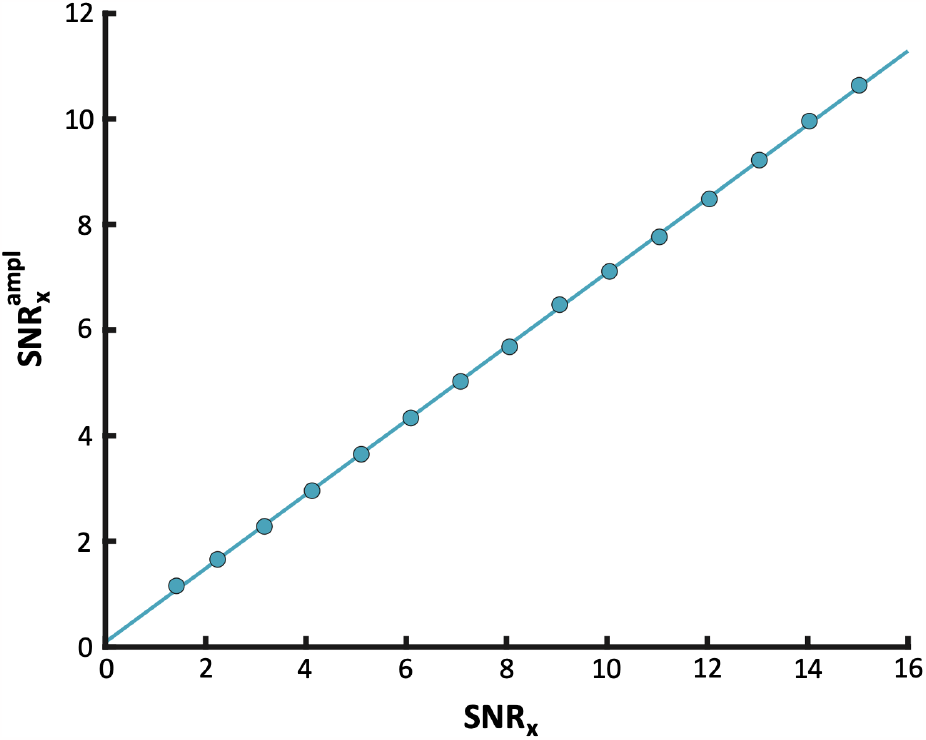
Linear relationship between the amplitude-specific SNR (Eq. 13) and the classical SNR (Eq. 4) for near-gaussian synthetic MEG signals (see main text). The regression model 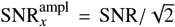 is superimposed (see last paragraph of this Appendix for the complete mathematical relationship).

The second term in Eq. (12) represents the spurious inflating effect of noise amplitude correlations on AEC, in complete analogy with the contribution of *R*_noise_ to linear correlation (Eq. 5). In principle, this term should vanish since a good model of “background noise” must be devoid of any amplitude coupling (see part i of noise model definition in section A.1), i.e., 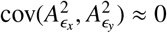. This was verified explicitly in the case of cerebral background noise modeled as non-bursting brain activity (see main text). So this effect only depends on noise linear correlations and is likely subdominant after leakage correction, but we nevertheless kept track of it for the sake of generality (see discussion in section A.1 below Eq. 6).

The third and last term involves the quantity

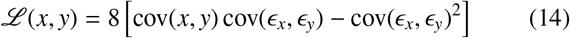

depending on the linear covariances between MEG signals *x, y* and between the corresponding noise signals *ϵ*_*x*_, *ϵ*_*y*_, along with a similar contribution *ℒ* (*x*, H*y*) that involves the Hilbert transforms H*y* and H*E*_*y*_ (see Eq. 18 below). This last contribution is proportional to the noise correlation *R*_noise_ and is thus likely subdominant after leakage correction. As we show below, *ℒ* (*x, y*) is also proportional to the neural zero phase-lag coupling *R*_0_ (and *ℒ* (*x*, H*y*) to the neural coupling at *π/*2 phase lag; see Eq. 28). Although *R*_0_ is generally thought to vanish in the resting state, Sjogard et al. (2019) suggested the existence of spontaneous quasi-zero lag correlations, at least within the default-mode network. For this reason, we will remain general and retain all contributions in Eq. (12).

Let us finally comment on the approximative nature of our result (12). It originates from two approximations needed to carry out analytical developments: Eq. (11) and deviations from the narrow-band limit (necessary to rely on the machinery of the Hilbert transform, see mathematical derivations below). Our validation with synthetic data (Fig. 1d) actually establishes the excellent accuracy of both approximations, so that for all practical purposes the approximation sign (≈) may thus be replaced by an equality in Eq. (12).

#### Power bias correction by renormalization

The general discussion that follows Eq. (5) holds without change here, and in particular the neural amplitude correlation AEC_0_ can be recovered via a renormalization procedure

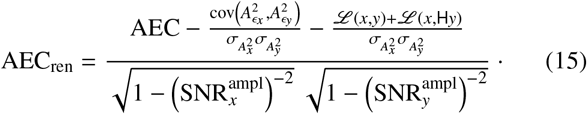

Critically, this quantity is expressed in terms of parameters that are experimentally accessible to MEG recordings. The two subtractive terms in the numerator correct for possible (and likely subdominant) noise correlations and zero phase-lag coupling, and the denominator corrects for the power bias itself. The estimation of second- and fourth-order statistics of the noise signals *ϵ*_*x*_ and *ϵ*_*y*_ relies once again on a separate modeling step of background noise. See the related discussion in Appendix A.1.

We now turn to the mathematical justification of our main theoretical result (12).

#### Mathematical derivations: Separation of timescales in oscillatory dynamics

We consider a general signal *x*(*t*), whose Fourier transform 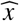 is supported on a frequency band of center *ν*_0_ and half-width Δ*ν < ν*_0_. Such signal can be factorized according to

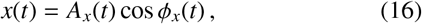

where the amplitude *A*_*x*_ and phase *ϕ*_*x*_ are determined through the relationship

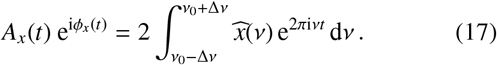

The Hilbert transform H*x* corresponds to the signal (16) but with 90-degree lagged phase *ϕ*_*x*_(*t*) − *π/*2, i.e.,

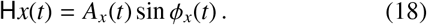

In the narrow-band limit Δ*ν* « *ν*_0_ pertaining to oscillatory dynamics, *A*_*x*_ captures slow modulations of the amplitude and *ϕ*_*x*_ rapid phase oscillations at an instantaneous frequency that varies slowly around the center frequency *ν*_0_. A succinct way to understand this basic property is to apply a change of variable *ν* → *ν*_0_ + *ν* in the Fourier integral (17), which yields

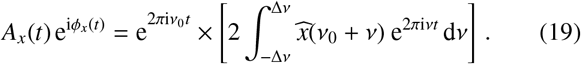

The factor between brackets in the right-hand side determines a complex-valued signal 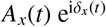 with same amplitude than *x*, a phase *δ*_*x*_ = *ϕ*_*x*_ − 2*πν*_0_*t* that corresponds to the phase shift between *x* and the carrying oscillation at frequency *ν*_0_, and a Fourier spectrum restricted to frequencies below Δ*ν*. This implies that the amplitude *A*_*x*_ and the phase shift *δ*_*x*_ both evolve on timescales much slower than the carrying oscillation.

#### Mathematical derivations: Technical identities on time averages

This separation of timescales applies to any narrow-band signal; in our setup *x* may be any one of the connectivity processes *x*_0_, *y*_0_ or background noises *ϵ*_*x*_, *ϵ*_*y*_. Together with the assumption that connectivity processes and noise are temporally independent (see noise model assumption ii in Appendix A.1), this is key to estimate analytically the several time averages that we will encounter below. The main results of this section are Eqs. (20), (24), and (29). They are presented here as approximate in the sense that they work in the narrow-band limit Δ*ν*« *ν*_0_, but this condition holds to a good approximation in our spectrally resolved MEG data (see main text) so in practice the approximation errors are small. Of note, no hypothesis on the statistical (in)dependence of amplitudes and phases will be required here.

While evaluating the correlation of amplitudes squared (11), a number of simplifications emerge from identities of the form

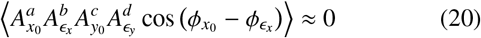

holding for any non-negative integers *a, b, c, d* ≥ 0, and similar identities involving the phase lag 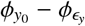. To demonstrate Eq. (20), we expand the left-hand side with a classical trigonometric identity and apply the independence of *x*_0_, *y*_0_ and *ϵ*_*x*_, *ϵ*_*y*_ to factorize time averages to obtain

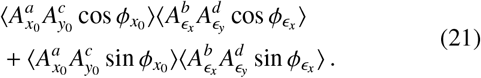

Each factor in this expression vanishes approximately because amplitudes remain approximately constant over any cycle of the phase oscillations. Let us consider for instance the first factor; inserting 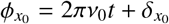 yields

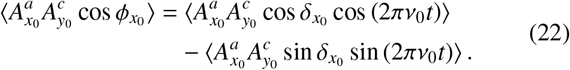

The Riemann-Lebesgue lemma establishes that each of these two terms vanish in the limit *ν*_0_*/*Δ*ν*→∞, but this formal argument can be explained on the basis of the separation of timescales. Slow amplitude and phase modulations involve frequencies below Δ*ν*« *ν*_0_ and so they are fairly constant over short time periods corresponding to one cycle of the rapid carrying oscillation at frequency *ν*_0_. This allows to factorize time averages into fast oscillatory averages over one cycle and slow modulatory averages over long times, e.g.,

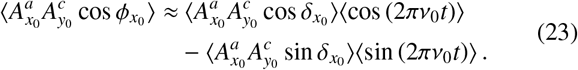

This expression vanishes since ⟨ cos (2*πν*_0_*t*) ⟩= ⟨ sin (2*πν*_0_*t*) ⟩ = 0 over any cycle. The same holds for all terms encountered above. This ends the demonstration of the identities (20).

We will also encounter non-vanishing averages involving second powers of cosines. First,

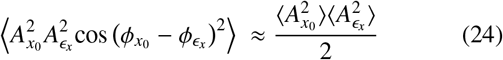

and likewise for the *y* signals. To derive Eq. (24), we again use trigonometry to rewrite the left-hand side as 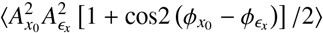. The cosine term vanishes in the narrow-band approximation for the exact same reasons than Eq. (20) holds, leaving 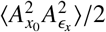. The latter average factorizes since connectivity processes and noise are independent, leading us back to Eq. (24).

The second non-vanishing identity is

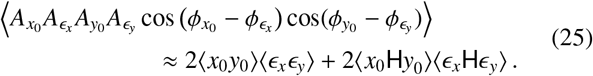

To establish this, we start by expanding both cosines as we did below Eq. (20). Recognizing the signals *x*_0_, *ϵ*_*x*_ and their Hilbert transform through Eqs. (16) and (18) allows to write

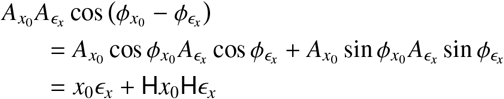

and similarly for the *y* signals. Inserting these relationships into the left-hand side of Eq. (25) yields a time average (*x*_0_*ϵ*_*x*_ + H*x*_0_H*ϵ*_*x*_) (*y*_0_*ϵ*_*y*_ + H*y*_0_H*ϵ*_*y*_). Expanding the product and using the independence of *x*_0_, *y*_0_ and *ϵ*_*x*_, *ϵ*_*y*_, we obtain

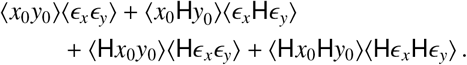

It turns out that the first and last terms are equal to each other, and likewise for the second and third terms, from which Eq. (25) follows. For example,

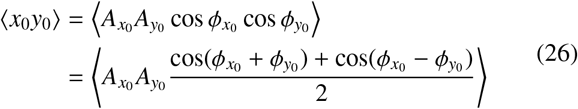

and

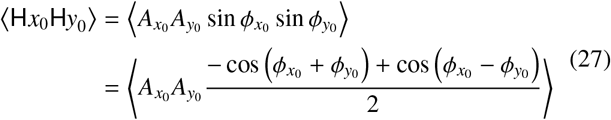

both converge to the same value 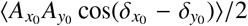 because the time average containing the high-frequency oscillation 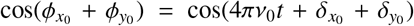 vanishes in the narrow-band approximation once again thanks to the separation of timescales.

It is noteworthy that the first term in the right-hand side of Eq. (25) is closely related to the quantity *ℒ* (*x, y*) defined in Eq. (14). More specifically,

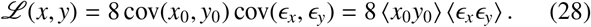

The first equality follows by inserting the relationship (2) between connectivity processes and measured signals into the definition (14) and using the independence of connectivity and noise processes to set both cov(*x*_0_, *ϵ*_*y*_) and cov(*ϵ*_*x*_, *y*_0_) to zero.

The second equality follows from the definition of covariance (reviewed right below Eq. 1) and the fact that all signal averages ⟨ *x*_0_⟩, ⟨ *y*_0_⟩, ⟨ *ϵ*_*x*_⟩, and ⟨ *ϵ*_*y*_⟩ vanish (which are particular cases of Eq. 20). Identity (25) can then be recast as

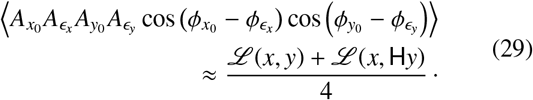

The significance of Eq. (28) is that it establishes the claim made below Eq. (14) that *ℒ* (*x, y*) is proportional to the neural zero-lag correlation *R*_0_. Similarly, by definition (18) of the Hilbert transform, *ℒ* (*x*, H*y*) is proportional to a measure corr[*x*_0_, H*y*_0_] of neural 90-degree phase lag coupling.

#### Mathematical derivations: Proof of Eq. (12)

After these technical preliminaries, we are now in position to derive our main theoretical result (12) from Eq. (11). Our starting point is the modulus squared of Eq. (10),

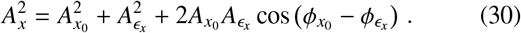

A similar relation holds for the signal *y*. These expressions play the same role here than Eq. (2) did in Appendix A.1, but the situation is somewhat more intricate due to the interplay between amplitude and phase-lag dynamics evidenced by the rightmost term in Eq. (30).

It is slightly simpler to deal with the denominator of Eq. (11) first. We insert the decomposition (30) into the defining expression of the variance of amplitude squared 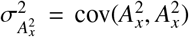. Expanding and gathering terms where possible, we find that 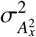 equals to

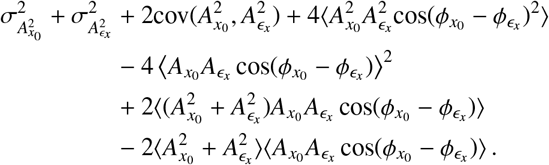

This daunting expression simplifies in the narrow-band limit, as the third term vanishes due to the independence of connectivity and noise processes (see model noise hypothesis ii in Appendix A.1) and so do the fifth to last terms thanks to identities (20). Our result (24) takes care of the fourth term and leads to

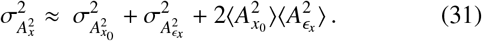

We then take the time average of the decomposition formula (30) and apply an identity (20) to simplify the last term,

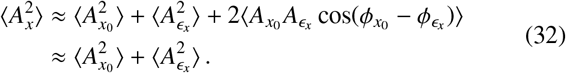

Combining the two preceding equations, we obtain

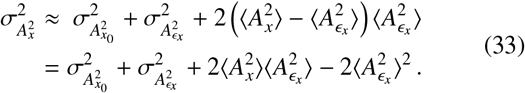

This is the analog of the intermediate result 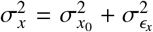 of Appendix A.1 but for amplitude squared. When placed in the denominator, we can recast Eq. (33) as

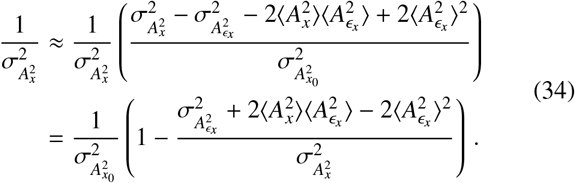

Comparison with the analogous result 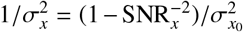 at the end of Appendix A.1 motivates our definition of the amplitude-specific SNR measure (13), since then

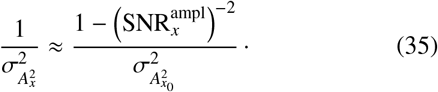

A similar result holds for the signal *y*. This handles the case of the denominator in Eq. (11); we now turn to its numerator.

We need to find the analog of the intermediate result cov(*x, y*) = cov (*x*_0_, *y*_0_) + cov(*ϵ*_*x*_, *ϵ*_*y*_) of Appendix A.1. Expanding the covariance 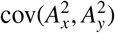 with the decomposition formula (30) and gathering terms leads to

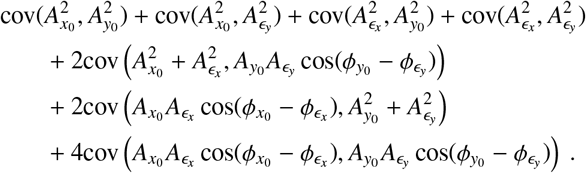

Again, several simplifications take place in the narrow-band limit. The second and third terms vanish according to the independence of connectivity and noise processes (see model noise hypothesis ii in Appendix A.1); the covariances in the fifth and sixth terms too since they involve averages of the form (20). The last term also simplifies to

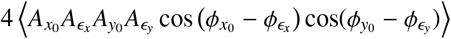

again thanks to Eq. (20), and thus it equals to *ℒ* (*x, y*) + *ℒ* (*x, Hy*) by our last identity (29). Gathering these observations together, we find

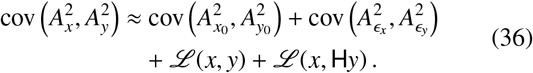

Our main result is now at hand. Equation (11) for the AEC consists in normalizing the covariance (36) of amplitudes squared by their standard deviation, which using Eq. (33) yields Eq. (12).

#### Mathematical derivations: Amplitude-specific SNR for gaussian signals

For completeness, we explore here the link between the amplitude-specific 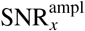 and the standard SNR_*x*_ in the case of gaussian signals, and thereby explain Fig. S1 analytically. We start from the observation that definition (13) involves a combination of averages of amplitude squared 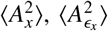 and to the fourth power 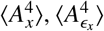, which can be expressed in terms of variance 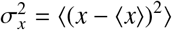 and fourth-order moment 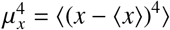 according to

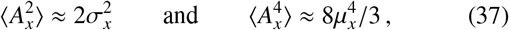

and similarly for *ϵ*_*x*_. These identities follow in the narrow-band limit from arguments very similar to above. Briefly, applying Eq. (16), the zero-mean property ⟨ *x*⟩ = 0, and trigonometric identities yields

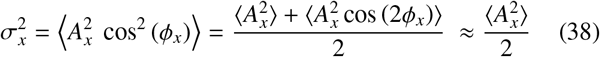

and

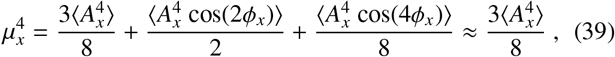

with the last equalities holding in the narrow-band limit.

Equations (37) allow to recast the definition (13) of the amplitude-specific SNR as

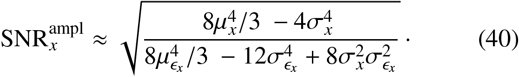

So far this result holds for arbitrary narrow-band signals, but it simplifies a great deal in the case of gaussian signals because they obey the zero-kurtosis identity 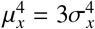. With the help of Eq. (4), we find

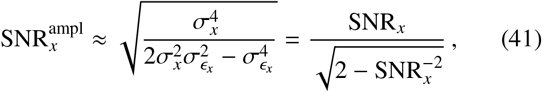

which comes close to 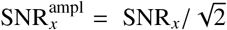 (except for SNR_*x*_ values very close to 1), in line with Fig. S1.

## Appendix B. SI Methods

### Appendix B.1. More on background noise models

Our procedure to model cerebral background noise allows to access noise statistics but not its full time course synchronized with experimental MEG signals (see properties i and ii in SI Theory, Appendix A.1). For example, coincident non-bursting activity only accounted for about one third of our recordings (average across subjects, connections, and frequencies) so it was not accessible two thirds of the time. One caveat with this abstract approach is that it does not guarantee that the resulting SNR estimates (4) lie above the theoretically minimal value 1 (set by Eq. 8), which would yield ill-conditioned or ill-defined renormalization (see discussion below Eq. 6 in SI Theory, Appendix A.1). This issue turned out to arise in a small fraction of our data, so in practice we conservatively excluded any connection for which the amplitude-specific SNR estimate (13) at any of its two nodes reached below 1.1 in at least one frequency band. This led to the exclusion of no more than 0.45% of our individual functional connectivity dataset (within a single subject) when modeling background noise as non-bursting activity, and 2% (spread over 6 out of the 31 subjects) when referring to measurement noise (i.e., empty-room MEG recordings nonsimultaneous to resting-state MEG recordings).

### Appendix B.2. More details on statistical procedures

#### Power bias measure (PBM)

We define here explicitly the PBM index in the slightly different cases considered in the main text. Each of them quantifies to what extent the power bias affects amplitude connectivity estimation by assessing the relative difference between the measured value AEC and either the ground truth AEC_0_ in simulations or the corrected value AEC_ren_ in resting-state data.

For the synthetic functional connectivity data, each of the *N*_subj_ = 30 “subjects” corresponded to a number of simulations where the true connectivity AEC_0_ was varied (simulating in a sense spectral variations of connectivity) while the SNR parameters (4) were kept fixed. The PBM was then taken as

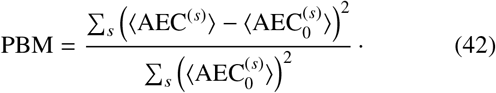

In this section, brackets ⟨ · ⟩denote group averaging over *N*_subj_ subjects (rather than time averaging as in SI Theory, Appendix A). Sums run here over all simulations (indexed by superscripts *s*) performed at fixed SNRs. This PBM index allowed us to quantify the effect of the power bias as a function of the SNR and was used in Fig. 1c; a large PBM indicates a strong effect of the power bias. A closely-related index was used to estimate the power bias correction error

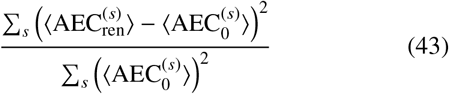

as a function of the SNR. A large value of this index indicates poor performance of the correction procedure (SI Theory, Appendix A.2, Eq. 15).

For the resting-state MEG connectivity data, the power bias was measured for each connection as

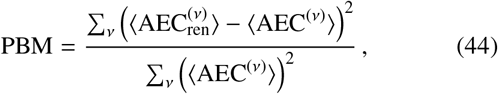

where AEC^(*ν*)^ corresponds to an individual amplitude correlation in the band of center frequency *ν*. Sums now run over all frequencies *ν* considered so as to quantify the effect of the power bias on group-averaged AEC spectra. Findings of large PBM demonstrate substantial modifications of connectivity values. This PBM is reported in the main text for interhemispheric network connectivity; a systematic analysis across the whole connectome is developed in SI Results (Appendix C.3). Of note, the square root of the denominator Σ_*ν*_(⟨ AEC^(*ν*^⟩^)^)^2^ in Eq. (44) corresponds to the “AEC spectrum norm” used to construct Fig. 2c.

#### Regularized Pearson correlation test of spectral similarities

Parametric correlation testing in power or connectivity spectra requires to know the number *n* of degrees of freedom in these spectra. We could not estimate it naïvely as the number *N*_freq_ = 45 of frequency bands because the natural bandwidth of neurophysiological processes leads to a degree of spectral smoothness (i.e., inter-dependent frequencies), so the resulting statistical test would be too lenient. Rather, we regularized our correlation tests in such a way that the null hypothesis adapts to the spectral smoothness, by setting *n* to an estimate of the number of effectively independent frequencies in the spectra under scrutiny. In practice, we computed the cross-frequency *N*_freq_ ×*N*_freq_ sample covariance matrix from individual spectra (averaged over the two spectra being correlated) and estimated *n* as the minimum number of eigenvalues summing up to 99% of the total variance.

### B.3. Statistical procedures for SI Results (Appendix C)

#### Relative contribution of power bias, noise correlations and linear synchronization

Given that the correction procedure controls for noise correlations and linear synchronization on top of the power bias (see SI Theory, Appendix A.2, Eq. 15), we further assessed the relative impact of the noise correlation term

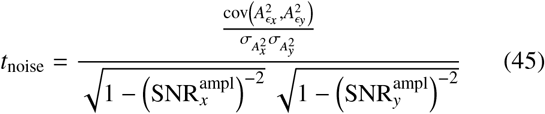

and of the synchronization term

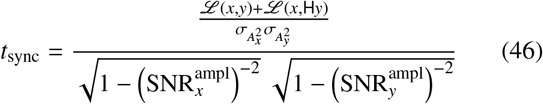

compared to the main power bias correction term

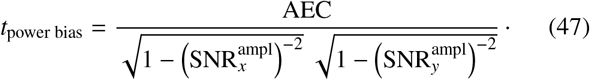

Each of these terms were estimated from resting-state MEG data separately for each subject, connection and frequency *ν*, and the relative contributions of noise correlations and linear synchronization were then measured as the squared spectrum norm ratios

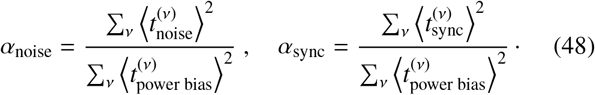

These relative contributions are reported in SI Results (Appendix C.1).

#### Multiple regression model for power bias correction

The power bias renormalization factor is nonlinear in the SNR (see SI Theory, Appendix A). To examine the importance of this feature, we considered another correction approach based on multiple regression. Regression models are widely used in neuroimaging to discard unwanted effects (covariates of no interest) and highlight effects thought to be relevant (covariates of interest). For each connection and each frequency band of our AEC data, we designed the linear model

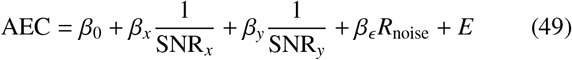

and determined the coefficients *β*_0_, *β*_*x*_, *β*_*y*_, and *β*_*ϵ*_ in a standard manner by minimizing the group variance of the model error *E*. To keep the approach generic and independent of the theoretical developments in SI Theory (Appendix A.2), we used the basic definition (4) of the SNR rather than the amplitude-specific form (Eq. 13) identified in the context of the power bias renormalization (Eq. 15), and likewise we used the linear noise correlation *R*_noise_. Nevertheless, we set the two SNR-related regressors to the inverse SNR as it is reasonable to expect on general grounds that connectivity estimates AEC converge to their true value AEC_0_ in the large SNR limit (SNR_*x*_»1 and SNR_*y*_» 1). The true value would then be captured by the intercept regressor corresponding to the first term of the above model. More specifically, *β*_0_ would correspond to the group-average of AEC_0_, and the error term *E* to the inter-individual variations.

The regression-based corrected amplitude connectivity estimate can thus be identified as

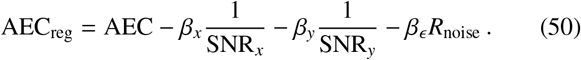

Again, this was applied independently for each pair *x, y* of source-projected MEG signals (with one being leakage corrected with respect to the other) and within each frequency band, and symmetrization was imposed afterwards (see Methods in the main text). The adequacy of this approach to power bias correction is examined in Fig. 3 for interhemispheric network connectivity and more systematically in SI Results (Appendix C.2).

#### Statistical assessment of connectivity changes: regularized Hotelling’s *T*^2^ test

The statistical significance of connectivity changes measured by the PBM (Appendix B.2) was established using a version of Hotelling’s *T* ^2^ test. Specifically, we computed for each connection the squared Mahalanobis distance in dimension *N*_freq_ = 45 between connectivity spectra AEC^(*ν*)^ before correction and 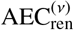 after correction (where *ν* denotes frequency), under the multivariate null hypothesis 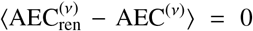 that the power bias does not affect AEC estimation. One issue in this computation is the non-invertibility of the corresponding *N*_freq_ × *N*_freq_ sample covariance matrix due to the inter-dependencies among frequencies (Appendix B.2). To regularize the situation, the covariance was pseudo-inverted using its first *n*′ largest eigenvalues, with *n*′ the number of effectively independent frequencies determined as in the case of the correlation tests (Appendix B.2) but applied here on the sample covariance of the differences between the two spectra being compared. Statistical inference was then performed at significance level *p <* 0.05 using Hotelling’s 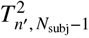 distribution in dimension *n*′ with *N* _subj_− 1 degrees of freedom. Of notice, *N*_subj_ was slightly smaller than the total number of subjects (i.e., 31) for a few, sparsely distributed connections due to the technical caveat mentioned in Appendix B.1. The results of this analysis are described in SI Results (Appendix C.3).

**Figure S2:**
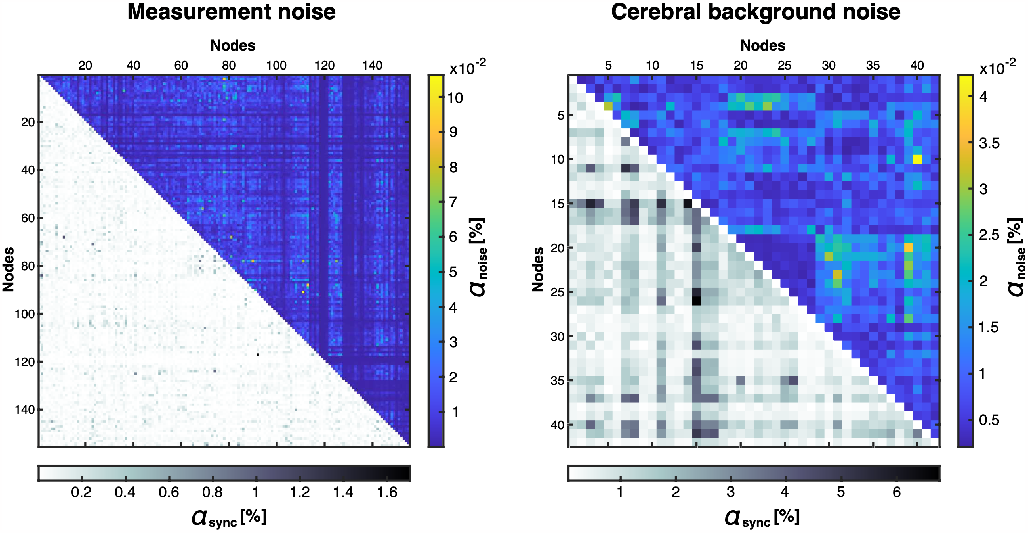
Relative contribution of correction features. The impact of noise correlation (*α*_noise_) and linear synchronization (*α*_sync_) in AEC power bias correction is mapped across the dense, 155-node connectome when using measurement noise (left) and across the sub-sampled, 42-node connectome when modeling cerebral background noise as coincident non-bursting activity (right). See SI Methods (Appendix B.3) for precise definitions of these percentages.

## Appendix C. SI Results

### Appendix C.1. Relative contribution of power bias, noise correlations and linear synchronization

Given that our correction method not only controls for the power bias but also for noise correlation and spontaneous linear synchronization (see SI Theory, Appendix A.2, Eq. 15), we assessed the contribution of the two latter factors relative to that of the power bias *per se* (Fig. S2). The contribution of noise correlations was extremely small compared to the power bias itself (*α*_noise_ *<* 0.11% when measurement noise is used in the correction, see Fig. S2, left; *α*_noise_ *<* 0.05% when cerebral background noise is modeled as non-bursting activity, see Fig. S2, right). This was expected by design of our approach to noise modeling since lack of connectivity was one defining property of noise (see SI Theory, Appendix A.1, property i) and in line with Fig. 2c, although in this case remnant noise correlations might subsist from possible spatial leakage miscorrections (Wens et al., 2015). The contribution of linear synchronization was larger, in line with the possible existence of spontaneous linear synchronization processes (Sjogard et al., 2019), but still highly subdominant compared to the power bias itself (*α*_sync_ *<* 1.7% when measurement noise is used in the correction, see Fig. S2, left; *α*_sync_ *<* 7% when cerebral background noise is modeled as non-bursting activity, see Fig. S2, right).

### Appendix C.2. Multiple regression model for power bias correction

To assess the importance of SNR nonlinearity in power bias correction, we considered linear regression of node SNRs (along with noise correlation) as an alternative correction approach. For conciseness, we focus here on power bias correction based on measurement noise where our nonlinear renormalization procedure (SI Theory, Appendix A.2, Eq. 15) led to virtually no connectivity changes (see main text and Appendix C.3); the results of regression modeling are thus quickly interpretable in this case.

**Figure S3:**
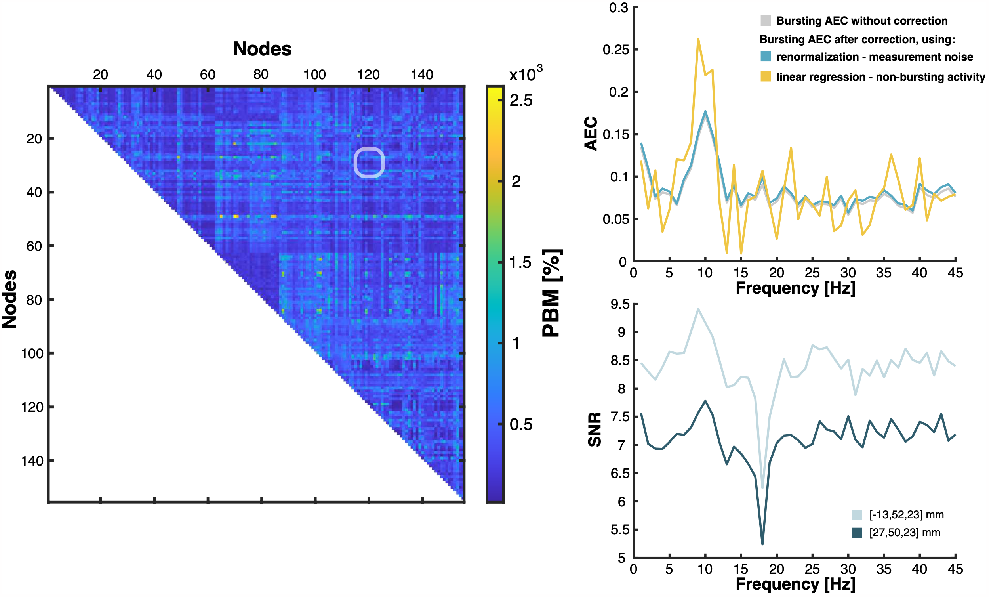
Power bias correction via linear regression in amplitude correlation spectra of the connectome. **Left**. The PBM (see SI Methods, Appendix B.2) associated with power bias correction based on regression and measurement noise (see SI Methods, Appendix B.3) was mapped in the whole-brain-covering connectome. **Right**. Functional connectivity (top) and SNR (bottom) spectra of the connection with the least amount of connectivity change. Its location in the connectome is highlighted in the corresponding PBM matrix.

The example of interhemispheric network connectivity considered in the main text shows that AEC spectra (Fig. 3, top, yellow) corrected by regression are much more erratic than renormalized AEC (Fig. 3, top, blue), especially in the AN. Accordingly, regression led to connectivity changes (SMN, PBM = 110%; AN, PBM = 550%; VN, PBM = 130%) hundreds of times larger than our renormalization (PBM = 1%, i.e., very small power bias effect size). This suggests that linear regression is inadequate for power bias correction, and that taking into account SNR nonlinearity is fundamental. This observation generalized across the whole connectome (Fig. S3).

We further sought to illustrate the reason for inadequacy of this regression graphically in Fig. S4, where we compare the nonlinear power bias model and the linear regression model from the SMN interhemispheric connectivity data at 9 Hz. The regression surface corresponds to a flat plane whose intercept value (i.e., the linear extrapolation of Eq. 49 in SI Methods, Appendix B.3 to very large SNR values 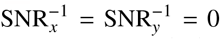 where power should not bias functional connectivity) yields the corrected AEC (Eq. 50). By contrast, the nonlinearity of our power bias model (Eq. 12 in SI Theory, Appendix A.2; see also Fig. 1b) yields a curved surface that flattens when approaching the intercept, which led to a lower renormalized AEC (Eq. 15). This nonlinear flattening of the power bias explains why the linear regression tends to overestimate interhemispheric AEC in Fig. 3.

### Appendix C.3. Statistical assessment of connectivity changes

In the main text, we focused on the question of whether or not power bias correction modifies the shape of connectivity spectra, not the global level of connectivity. Here we provide a quantitative and statistical analysis of the sheer effect size of connectivity changes brought by the power bias and measured by the PBM index.

**Figure S4:**
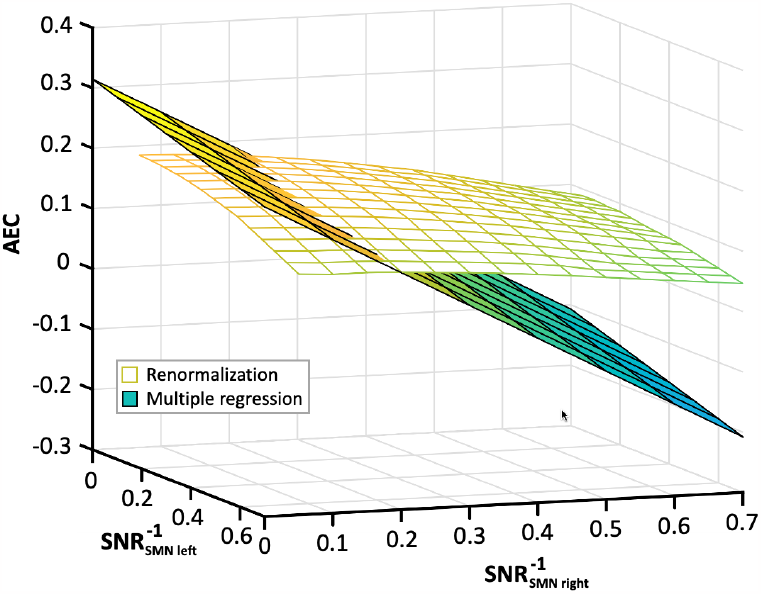
Comparison of power bias correction and multiple regression, using measurement noise. This example taken from the resting-state SMN connectivity data at 9 Hz shows two model surfaces used for two distinct correction methods. The planar surface corresponds to multiple regression and the curved one, to power bias correction by AEC renormalization (15). In both cases, the correction consists in extrapolating AEC to the intercept value above 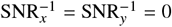.

**Figure S5:**
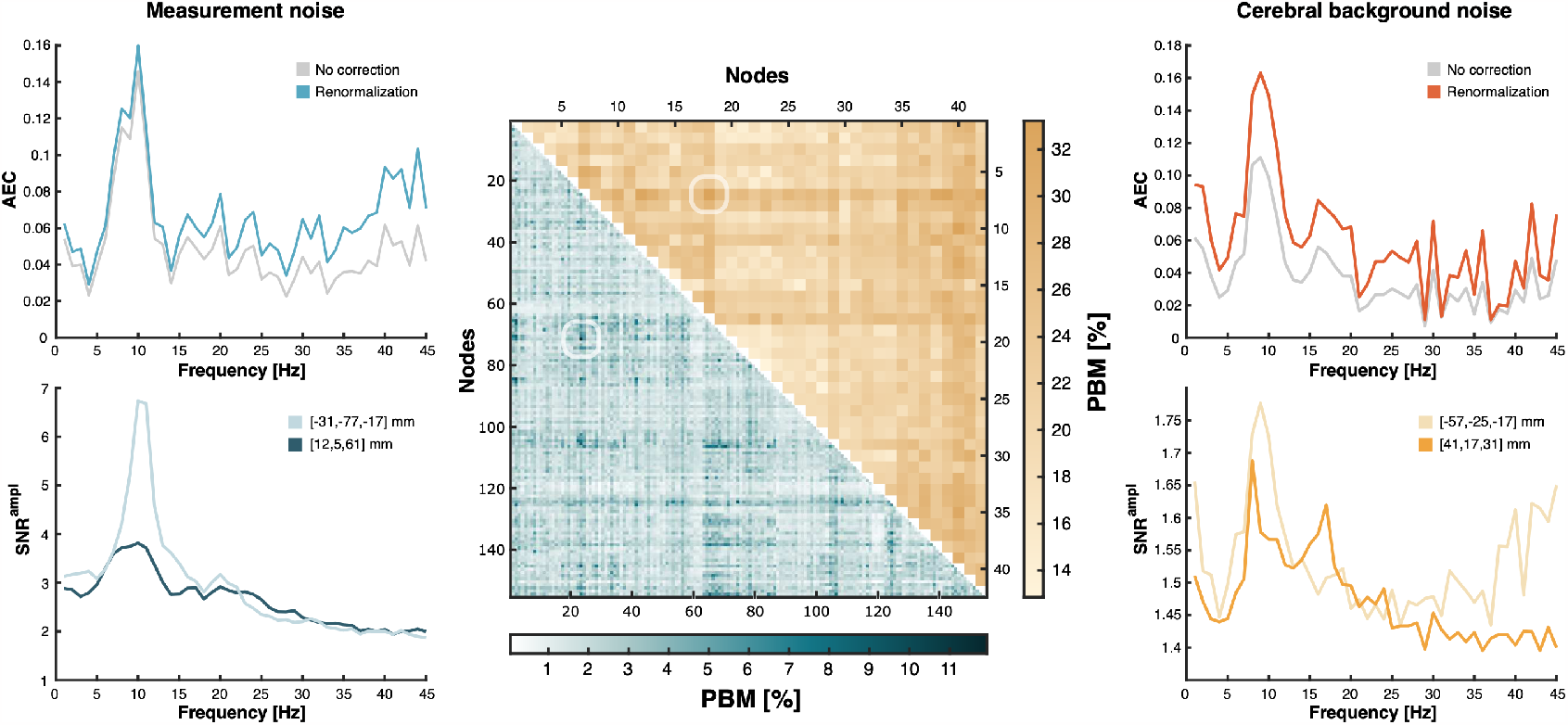
Power bias effect size in amplitude correlation spectra of the connectome. **Middle**. The PBM (see SI Methods, Appendix B.2) associated with power bias correction relative to measurement noise (lower-left triangle, blue shades) or to cerebral background noise modeled as non-bursting activity (upper-right triangle, yellow shades) was mapped in the whole-brain-covering connectome. **Left**. Functional connectivity (top) and SNR (bottom) spectra of the connection showing the highest connectivity change when modeling noise as measurement noise. Its location in the connectome is highlighted in the corresponding PBM matrix. **Right**. Same as left, but modeling cerebral background noise as non-bursting brain activity.

#### Interhemispheric connectivity

When modeling cerebral background noise as non-bursting brain activity, the power bias levelled up interhemispheric network connectivity by about 20% (PBM values; SMN, 18%; AN, 26%; VN, 17%), but these effects were not statistically significant (regularized *T* ^2^ test; SMN, *p* = 0.065; AN, *p* = 0.27; VN, *p* = 0.051). The PBM dropped substantially (to 1% for the three networks) when using measurement noise, yet rather counter-intuitively these small effects turned out marginally significant (regularized *T* ^2^ test; SMN, *p* = 0.007; AN, *p* = 0.053; VN, *p* = 0.025). This was explained by the observation that our cerebral background noise model based on non-bursting brain states revealed a higher inter-individual variability in corrected functional connectivity (that we presume of biological origin) than measurement noise (which corresponds to highly reproducible empty-room MEG recordings).

#### Functional connectome

Figure S5 reports a systematic analysis of PBM across the connectome. The strongest effect sizes were located at a connection linking the ventral-attentional (node MNI coordinates, [41, 17, 31] mm) and the default-mode networks ([−57, −25, −17] mm) with a PBM = 33% change for power bias correction relative to cerebral background noise (Fig. S5, right), and at a connection linking the ventral-attentional ([12, 5, 61] mm) and the visual networks ([−31, −77, −17] mm) with a PBM = 12% change for power bias correction relative to measurement noise (Fig. S5, left). However, none of these connectivity changes turned out statistically significant after controlling for the familywise error rate.

